# Towards HCP-Style Macaque Connectomes: 24-Channel 3T Multi-Array Coil, MRI Sequences and Preprocessing

**DOI:** 10.1101/602979

**Authors:** Joonas A. Autio, Matthew F. Glasser, Takayuki Ose, Chad J. Donahue, Matteo Bastiani, Masahiro Ohno, Yoshihiko Kawabata, Yuta Urushibata, Katsutoshi Murata, Kantaro Nishigori, Masataka Yamaguchi, Yuki Hori, Atsushi Yoshida, Yasuhiro Go, Timothy S. Coalson, Saad Jbabdi, Stamatios N. Sotiropoulos, Stephen Smith, David C. Van Essen, Takuya Hayashi

## Abstract

Macaque monkeys are an important model species for understanding cortical organization of primates, yet tools and methods for noninvasive image acquisition (e.g. MRI RF coils and pulse sequence protocols) and image data preprocessing have lagged behind those developed for humans. To resolve the structural and functional characteristics of the relatively thin macaque cortex, high spatial, temporal, and angular resolutions are required while maintaining high signal-to-noise ratio to ensure good image quality. To address these challenges, we developed a macaque 24-channel receive coil for 3-T MRI with parallel imaging capabilities. This coil enabled adaptation of the Human Connectome Project (HCP) image acquisition protocols to the macaque brain. We also adapted HCP preprocessing methods optimized for the macaque brain, including spatial minimal preprocessing of structural, functional MRI (fMRI), and diffusion MRI (dMRI). The coil provided high signal-to-noise ratio and high efficiency in data acquisition, allowing four- and five-fold acceleration for dMRI and fMRI, respectively. Automated parcellation of cortex, reconstruction of cortical surface, removal of artefacts and nuisance signals in fMRI, and distortion correction of dMRI performed well, and the overall quality of basic neurobiological measures was comparable with those for the HCP. The resulting HCP-style in vivo macaque MRI data show considerable promise for analyzing cortical architecture and functional and structural connectivity using advanced methods that have previously only been available for humans.

**Highlights:**

➢ 24-channel 3T MR receive coil designed for the smaller macaque brain.
➢ In vivo macaque imaging protocols adapted according to guidelines from the HCP.
➢ Parallel imaging yields five- and four-fold acceleration in fMRI and dMRI sampling.
➢ HCP’s minimal preprocessing and denoising pipelines adapted for macaques.
➢ The multi-modal MRI data show considerable promise for HCP-style analyses.

## Introduction

Old World monkeys are an important neuroscientific model for understanding primate neuroanatomy (Brodmann K., 1905; Felleman and Van Essen, 1991; Van Essen et al., 2001). Macaque monkeys have provided insights about neurovascular coupling (Goense and Logothetis, 2008), neural wiring (Markov et al., 2014) and the evolution of the human brain’s functional connectome (Passingham, 2009; Wang et al., 2012). However, macaques are separated from humans by 25 million years of evolution, and are known to have substantial brain differences despite being members of the same primate order. Recent imaging studies have revealed substantial neuroanatomical differences between macaques and humans, for example in language connectivity or proportion of cortex devoted to lightly myelinated association areas (Donahue et al., 2018; Glasser et al., 2014; Rilling et al., 2008). At the level of cortical areas, high confidence homologies (i.e., a common evolutionary origin) have only been firmly established for a modest number of early sensory and motor areas (Van Essen and Dierker, 2007) but are more challenging to delineate for higher cognitive regions such as prefrontal cortex (Mars et al., 2018b, 2018a). Improvements to in in vivo neuroimaging acquisition and preprocessing may help address several outstanding questions: what is the optimal interspecies registration between macaque and human cerebral cortices? What are the optimal methods for non-invasively estimating functional and structural connectivity as assessed by comparison with gold standard invasive tracers in macaques? What brain networks are shared and which ones are different between macaques and humans?

Recently, the Human Connectome Project (HCP) developed an improved, integrated approach to brain imaging acquisition, analysis, and data sharing (Glasser et al., 2016b). The overall goal of this approach is to increase the sensitivity and precision with which brain imaging studies are conducted in the hope that this will yield results that are more neurobiologically interpretable and more accurately comparable across individuals and studies. The HCP-style approach has seven core tenets (Glasser et al., 2016b): 1) Acquire as much high-quality data from as many subjects as possible. 2) Acquire data with maximum feasible resolution in space and time 3) Preserve high data quality throughout preprocessing by removing physical distortions, subject movement within and between scans, image intensity inhomogeneities, and artefacts and nuisance signals without blurring the data or altering the neural signals (Andersson et al., 2003; Andersson and Sotiropoulos, 2016; Glasser et al., 2013, 2016b, 2017; Griffanti et al., 2014; Salimi-Khorshidi et al., 2014). 4) Use appropriate geometrical models—surfaces for the sheet-like cerebral cortex and volumes for globular subcortical structures (Glasser et al., 2013). 5) Align brain areas across subjects, not cortical folds (Robinson et al., 2018, 2014). 6) Use a data-driven structurally and functionally sensible parcellation, ideally derived from multiple modalities (Glasser et al., 2016a). 7) Share results as data files in neuroimaging databases such as the Brain Analysis Library of Spatial maps and Atlases (BALSA) database (Van Essen et al., 2017), not just 3D coordinates. Following the HCP-Style approach leads to dramatic improvements in spatial localization precision in humans relative to traditional brain imaging methods (Coalson et al., 2018). Therefore, we sought to bring this improved brain imaging approach to non-human primate studies.

Monkey brains present distinct imaging-related challenges relative to human brains. The macaque brain is 10-fold smaller in weight, and its neocortex is ~25% thinner (average 2.0 mm vs 2.6 mm; (Donahue et al., 2018; Glasser et al., 2016b)). These facts necessitate increased spatial resolution to achieve comparable neuroanatomical resolution; however, smaller voxels are associated with decreased signal-to-noise ratio (SNR). One way to improve SNR is to scan at ultrahigh magnetic field strength (e.g., 7T). However, 7T scanners are not widely available and in any event pose technical challenges such as increased B_0_ and B_1_ inhomogeneity (Van de Moortele et al., 2009). For conventional 3T scanners, one key factor to enable high-resolution whole-brain imaging in macaques is to optimize the multi-channel radiofrequency (RF) receiver coil. Using a coil matched to macaque head size with a large number of small coil elements can yield improvements in SNR. Multi-channel signal acquisition using advanced 3T research scanners in humans enables parallel imaging both in the slice direction (i.e. multiband) (Moeller et al., 2010; Setsompop et al., 2012) and within the slice plane (generalized auto-calibrating partially parallel acquisitions [GRAPPA]) (Griswold et al., 2002). Although several studies have devised multichannel receive coils for macaque whole-brain imaging at 3T (Helms et al., 2013; Janssens et al., 2013, 2013; Khachaturian, 2010) and 7T (Gilbert et al., 2016; Mareyam et al., 1823), they have not to date demonstrated robust whole-brain mapping of multi-modal MRI measures such as those acquired by HCP. Achieving comparable results in macaques requires not only higher resolution and SNR but also low geometric distortion and signal intensity inhomogeneity, and requires optimized hardware, sequences, and post-processing techniques.

In this study, we designed and built a 24 channel receive coil with a geometry optimized for parallel imaging of anesthetized macaque monkeys at 3T. Capitalizing on the accelerated imaging capabilities of the coil, we adapted HCP-style data acquisition protocols for structural MRI (Glasser et al., 2013), fMRI (Smith et al., 2013) and diffusion MRI (dMRI) (Sotiropoulos et al., 2013; Uğurbil et al., 2013) to the small size of the macaque brain, as well as the HCP-style minimal spatial preprocessing and denoising pipelines (Andersson and Sotiropoulos, 2016; Glasser et al., 2018, 2016a, 2013; Salimi-Khorshidi et al., 2014, 2014). We generate accurate white and pial cortical surfaces, subcortical segmentations, myelin maps, and cortical thickness maps from structural MRI, surface aligned fMRI dense timeseries that have spatial artefacts and nuisance signals removed, resting state functional networks, and diffusion-based fiber orientation estimates, example tractography connections, and cortical neurite orientation and dispersion imaging (NODDI) (Zhang et al., 2012). The spatial resolution of the structural and functional imaging modalities are scaled to the macaque cortical thickness, thus providing comparable neuroanatomical resolution to HCP-style human imaging and facilitating comparison of connectomes between macaques and humans.

## Methods and Materials

Experiments were performed using a 3T clinical MRI scanner (MAGNETOM Prisma, Siemens, Erlangen, Germany) equipped with 80 mT/m gradients (XR 80/200 gradient system with slew rate 200 T/m/s) and a 2-channel B_1_ transmit array (TimTX TrueForm). The animal experiments were conducted in accordance with the institutional guidelines for animal experiments and animals were maintained and handled in accordance with the Guide for the Care and Use of Laboratory Animals of the Institute of Laboratory Animal Resources (ILAR; Washington, DC, USA). All animal procedures were approved by the Animal Care and Use Committee of the Kobe Institute of Riken (MA2008-03-11). We also used HCP data as a reference for data quality. The use of HCP data was approved by the institutional ethical committee (KOBE-IRB-16-24).

### Macaque 24-channel coil

The coil frame geometry was designed using a 3D digital design software (Rhinoceros 5, McNeel, Seattle, USA) to closely fit the head geometry of the animal with largest head dimensions (anterior posterior 109 mm, left-right 99 mm, superior-inferior 84 mm) (Fig. 1A) in our macaque head MRI database. The database included structural scans of 133 individual subjects from three macaque species (*Macaca fuscata*, N=4; *Macaca fascicularis*, N=122; *Macaca mulatta*, N=7). The largest animal’s MRI data was used to delineate the contour of the head surface and imported into the 3D digital design software where the inner surface of the coil was designed to closely fit the surface of the head (Fig. 1A). Next, 16 pentagonal and 8 hexagonal elements were configured over the surface (Fig. 1B), resembling a soccer-ball coil design (Wiggins et al., 2006). These elements were arranged in three quasi-horizontal arrays to maximize parallel encoding power of multiband EPI sequences for animals placed in the supine position and axial slices. The inner body of the device was constructed using a 3D printer (M200, Zortrax, Olsztyn, Poland) (Fig. 1C), and the coil elements were arranged over its external surface. Initially the coil elements were wired using a thin copper foil-plate (width 5 mm); however, because the plate elements markedly interfered with B_1_ transmission (data not shown), the coil elements were rewired using thin coaxial copper cables (Fig. 1D, cable diameter 0.7 mm; cable loop maximum mean diameter 48.6 ± 8.7 mm) (Wiggins et al., 2009), which substantially reduced interference with B_1_ transmission. The elements were arranged to continuously (critically) overlap each other to reduce coupling between nearest-neighbor coils (Roemer et al., 1990), and those in the caudal-posterior part were designed to have relatively larger diameter (35% larger in maximum diameter) to increase sensitivity to distant brain regions (e.g. cerebellum) while reducing sensitivity to closer regions (e.g. occipital cortex). The two elements placed over the eyes were also relatively large in diameter to allow video recording of eyes and eyelids for monitoring depth of anesthesia. In addition, capacitors were arranged vertically against the surface of the coil frame to reduce interaction with B_1_-transmission (Fig. 1D). Fig. 1E shows the circuits, which followed a standard design (Wiggins et al., 2006) consisting of diode detuning trap, cable trap and bias T connected to low input-impedance preamplifiers (Siemens Healthcare, Erlangen, Germany). The completed coil is shown in Fig. 1F.

**Figure 1.**
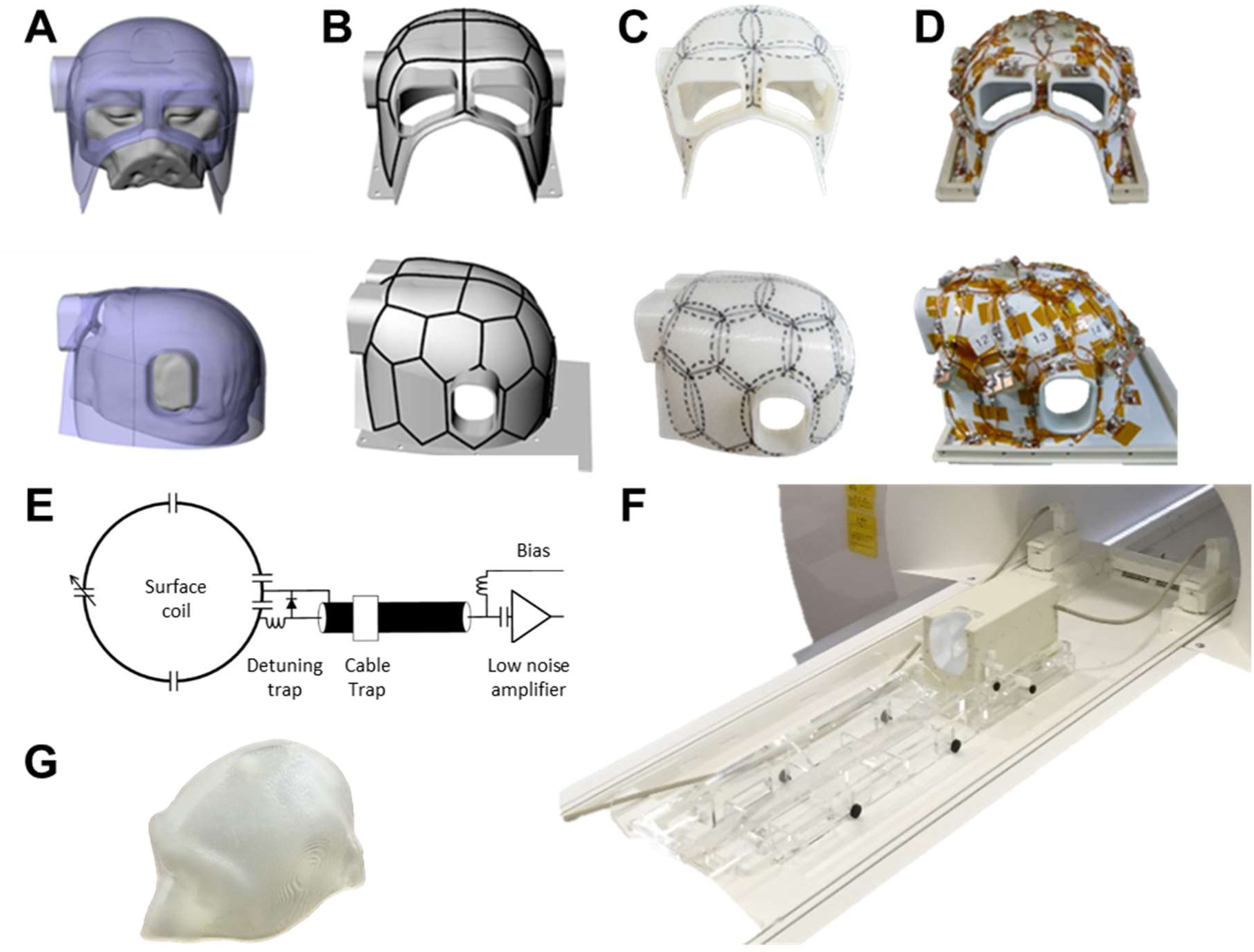
The design and development of macaque 24-channel receive-only coil. (**A**) Design of coil geometry and **(B)** element locations. **(C)** Outlook of element alignment on a 3D print. **(D)** Coil with final element arrangements. **(D)** Schematic of a coil element circuit. (**E**) Coil circuitry. **(F)** Coil outlook with animal holder attached to the gantry of the MRI scanner. **(G)** Macaque head phantom.

### Coil Evaluation

Coil elements were assessed for the ratio of loaded to unloaded quality factor Q, nearest-neighbor coupling, and active detuning. Element coupling was also estimated with gradient off-line noise correlation measurements. Two phantoms (NaCl 0.9%, gadolinium 0.1 mM) were designed and prepared using a 3D printer: one to closely match the inner-surface of the coil (Fig. 1G) used for B_1_ quality evaluation and the other to match to a typical macaque brain size used for geometry-dependent noise amplification. B_1_-transmission was assessed with a vendor provided flip-angle sequence. B_1_-receive field was estimated using a gradient-echo sequence and by calculating the signal ratio between 24-channel and body receive coils. Finally, geometry-dependent noise amplification due to parallel imaging was evaluated using gradient-echo imaging and GeneRalized Autocalibrating Partial Parallel Acquisition (GRAPPA) (Griswold et al., 2002) in-plane acceleration factors of 2, 3 and 4.

### Data Acquisition Strategy – Resolution and Contrast Considerations

To improve comparability of macaque and human brains, our data acquisition strategy sought to obtain data following methodologies introduced by the HCP (3T protocols) (Glasser et al., 2016b, 2013; Smith et al., 2013; Sotiropoulos et al., 2013; Uğurbil et al., 2013). To accurately model the cortical pial and white matter surfaces, structural imaging spatial resolution target (0.5 mm isotropic in macaques, equivalent to 0.8mm in humans) was based on preliminary evaluations of macaque cortical thickness (Glasser et al., 2014) and corresponds to approximately half of the minimum cortical thickness in the cortex, which is ≈1 mm in macaques (Donahue et al., 2018) and 1.6 mm in humans (Glasser et al., 2016b). Tissue contrast (grey and white matter and CSF) associated imaging parameters (e.g. inversion time, flip angle, repetition time and echo-time) were experimentally adjusted to produce robust surface estimation within the FreeSurfer pipelines, in conjunction with maximizing intracortical T1w/T2w (myelin-related) contrast. The fMRI spatial resolution selection (1.25 mm) was based on preliminary evaluations of the 5^th^ (low) percentile of cortical thickness, a similar strategy in humans by the HCP (resolution of 2 mm) (Glasser et al., 2016b). The temporal sampling rate (TR=0.75 sec) was maximized according to tSNR (see below) which was close to the human protocol (0.72 sec; Smith et al., 2013). For dMRI, the smallest spatial resolution within the practical limitation of the SNR was chosen using the same b-value scheme (b = 1000, 2000 and 3000 s/mm^2^) as in the HCP (Sotiropoulos et al., 2013) with 500 directions (more than the 270 in the HCP). Pilot studies for each modality included assessments for varying spatial resolution, flip-angle, RF transmission power, pulse length, inversion time (TI), fat suppression, multiband acceleration factor, in-plane acceleration factor, repetition time (TR), echo-time (TE), echo-spacing, spectral width, phase encoding direction, phase partial Fourier, phase oversampling, image resolution and diffusion directions.

### Structural Acquisition Protocol

T1w images were acquired using a 3D Magnetization Prepared Rapid Acquisition Gradient Echo (MPRAGE) (Mugler and Brookeman, 1990) sequence (0.5 mm isotropic, FOV=128×128×112 mm, matrix=256×256 slices per slab=224, coronal orientation, readout direction of inferior (I) to superior (S), phase oversampling=15%, averages=3, TR=2200 ms, TE=2.2 ms, TI=900 ms, flip-angle=8.3°, bandwidth=270 Hz/pixel, no fat suppression, GRAPPA=2, turbo factor=224 and pre-scan normalization). The value of TI (900 ms) was selected based on the contrast between white and grey matter and SNR. T2w images were acquired using a Sampling Perfection with Application optimized Contrast using different angle Evolutions (SPACE) sequence (Mugler et al., 2000) (0.5 mm isotropic, FOV=128×128×112mm, matrix=256×256, slice per slab=224, coronal orientation, readout direction I to S, phase oversampling=15%, TR=3200 ms, TE=562 ms, bandwidth=723 Hz/pixel, no fat suppression, GRAPPA=2, turbo factor=314, echo train length=1201 ms and pre-scan normalization). The total acquisition time for structural scans was 22 min (17 min for T1w and 5 min for T2w).

### Functional Acquisition Protocol

To reduce susceptibility induced geometric distortions and signal loss, the data was acquired in LR and RL directions. Functional scans were acquired using gradient-echo EPI (FOV=95×95 mm, matrix=76×76, 1.25 mm isotropic, interleaved slice order, and number of slices=50 covering the whole brain).

An empirical estimate of the effect of multiband slice acceleration factor on fMRI tSNR was obtained by a procedure similar to that used by the HCP (Smith et al., 2013). Briefly, simultaneous slice excitation enables a multiband factor fold reduction in the TR and subsequent incomplete T1-recovery leads to a reduction in the optimal (Ernst) flip angle and thus in tSNR. However, as more data volumes can be acquired in a matched time window, a more relevant estimate for the data quality can be calculated by multiplying the tSNR with a square root of acquired data timepoints. Therefore, tSNR was estimated with a matched image acquisition time (10 min) using a range of multiband factors (1, 3, 5, 6 and 8), minimum excitation and refocus RF-pulse lengths (with constant spectral width), TRs (3850, 1300, 840, 680 and 530 ms), corresponding (blood) Ernst angles (86, 65, 55, 51 and 45°) and a fixed bandwidth (1370 Hz/pixel).

These trials led us to select the imaging parameters (multiband factor=5, TR=755 ms, number of slices=45, flip-angle=55°, TE=30 ms, bandwidth=1370 Hz/pixel and echo spacing=0.95 ms and pre-scan normalization) for the fMRI data acquisition. To maintain the temporal autocorrelation structure of the data, long continuous runs were used (single-run scan time 51 min, 4096 frames, RL and LR directions resulting in a total acquisition time of 102 min).

### Field-Map Acquisition Protocol

The B_0_ field-map was estimated using a pair of spin-echo EPI images with opposite phase encoding directions (Andersson et al., 2003) (LR and RL directions, FOV=95×95 mm, 1.25 mm isotropic resolution, axial orientation, slices=45, interleaved data acquisition, TE=46.2 ms, 6/8 phase partial Fourier, bandwidth=1370 Hz/pixel, echo spacing=0.95 ms, fat suppression and pre-scan normalization). The B_1_ transmit field-map was obtained using vendor provided flip-angle sequence (Siemens, B_1_-map) (FOV=128×128×58mm, gap=2 mm, gaps acquired in a separate run, 2 mm isotropic, TR=10 s, target flip-angle=90°).

### Diffusion Acquisition Protocol

Diffusion scans were acquired with a 2D spin-echo EPI Stejskal-Tanner sequence (Stejskal and Tanner, 1965), utilizing monopolar gradient scheme and gradient pre-emphasis to reduce eddy currents. The monopolar gradients allowed decreased TE and significantly improved SNR without significant degradation due to eddy currents (in part due to the correction for eddy currents in post processing) (Andersson et al., 2003). The diffusion scheme contained three shells with b-values of 1000, 2000 and 3000 s/mm^2^ (diffusion time=26.5 ms, gradient duration=17.8 ms and amplitude=69.7 T/m),in accordance with the HCP (Sotiropoulos et al., 2013), but the number of direction (N_D_) was increased to 500 uniformly distributed over the sphere, as compared with that in the HCP (N_D_=270). Furthermore, 52 b=0 s/mm^2^ volumes were evenly distributed across the diffusion scheme to reduce CSF pulsation related uncertainty in the b=0 s/mm^2^ image signal intensity. In contrast to the HCP, we used GRAPPA (acceleration factor= 2) to reduce image distortions and accelerate the sequence in plane with a more recent version of the multiband sequence than was available for the original young adult HCP (Uğurbil et al., 2013). To correct for geometric distortions, the diffusion scheme was obtained using two scans with reversed phase encoding directions (LR and RL) and different number and directions of diffusion gradient (252 and 248) (Andersson and Sotiropoulos, 2016). The following imaging parameters were applied: FOV=90 mm, matrix=100×100, 0.9 mm isotropic resolution, number of slices=60, interleaved slice acquisition, multiband factor=2, GRAPPA= 2, TR=3400 ms, flip-angle=90, TE=73 ms, 6/8 phase partial Fourier, echo spacing=1.12 ms, bandwidth=1086 Hz/pixel, pre-scan normalization on and fat suppression using gradient reversal technique (Gomori et al., 1988). Total acquisition time was 30 min, during which frequency drift was small (≈0.5 Hz/min). By applying slice and in-plane accelerations (2×2), the acquisition time was reduced by more than 3-fold than without acceleration. However, the shortest possible TR was not used, in order to preserve SNR (to allow near-complete longitudinal magnetization recovery).

### Animal experiments

Macaque monkeys (mean 5380 g, range 3030–8850 g) were initially sedated with intramuscular injection of dexmedetomidine (4.5 μg/kg) and ketamine (6 mg/kg). A catheter was inserted into the caudal artery for blood-gas sampling, and tracheal intubation was performed for steady controlled ventilation using an anesthetic ventilator (Cato, Drager, Germany). End-tidal carbon dioxide was monitored and used to adjust ventilation rate (0.2 to 0.3 Hz) and end-tidal volume. After the animal was fixed in an animal holder, anesthesia was maintained using intravenous dexmedetomidine (4.5 μg/kg/hr) and 0.6 % isoflurane via a calibrated vaporizer with a mixture of air 0.75 L/min and O_2_ 0.1 L/min. Rectal temperature (1030, SA Instruments Inc., NY, USA) and peripheral oxygen saturation and heart rate (7500FO, NONIN Medical Inc., MN, USA) were monitored throughout experiments. For diffusion imaging the level of isoflurane was increased to 1.0 % to reduce potential eye and head motion artefacts.

### Data analysis

Data analysis utilized a version of the HCP pipelines with some customized specifically for use with non-human primates including structural (PreFreeSurfer, FreeSurferNHP (instead of FreeSurfer) and PostFreeSurfer), functional (fMRIVolume, fMRISurface) and diffusion preprocessing (DiffusionPreprocessing) (Donahue et al., 2016; Glasser et al., 2013). These NHPHCP pipelines requires FMRB’s Software Library (FSL) v6.0.1, FreeSurfer v5.3.0-HCP and Connectome Workbench v1.3.2 (https://www.humanconnectome.org/software/get-connectome-workbench) and are available at https://github.com/Washington-University/NHPPipelines.

### Structural Image Processing

Preprocessing began with the PreFreeSurfer pipeline, in which structural T1w and T2w images were registered into an anterior-posterior commissural (AC-PC) alignment using a rigid body transformation, non-brain structures were removed, T2w and T1w images were aligned using boundary based registration (Greve and Fischl, 2009), and corrected for signal intensity inhomogeneity using B_1_-bias field estimate. Next, data was transformed into a standard “Yerkes19” macaque atlas (Donahue et al., 2018, 2016) by 12-parameter affine and nonlinear volume registration using FLIRT and FNIRT FSL tools (Jenkinson et al., 2002).

Then, the FreeSurferNHP pipeline reconstructed the cortical surfaces using FreeSurfer v5.3.0-HCP (Fischl, 2012). This process included conversion of data in AC-PC space to a ‘fake’ space with 1-mm isotropic resolution in volume with a matrix of 256 in all directions, intensity correction, segmentation of the brain into cortex and subcortical structures, reconstruction of the white and pial surfaces and estimation of cortical folding maps and thickness. The intensity correction was performed using FMRIB’s Automated Segmentation Tool (FAST) (Zhang et al., 2001) followed by scaling the whole brain intensity by a species-specific factor (=80). This process significantly improved white and grey contrast particularly in the anterior temporal lobe as well as white surface estimation, an effect that may be associated with the so-called ‘anterior temporal lobe problem’ in pediatric brains, potentially due to less myelination in these white matter areas. We also improved the subcortical parcellation training dataset for the macaque brain, and trained for 21 subcortical structures: brainstem plus bilateral accumbens, amygdala, caudate, claustrum (which is not a part of the default structures for human FreeSurfer), cerebellum, diencephalon, hippocampus, pallidum, putamen, and thalamus (Fischl et al., 2002). The training dataset for brain mask extraction was also created. After parcellating the cortical and subcortical structures with these training datasets using the T1w image, the claustrum was treated as putamen, so that subsequent white surface estimation accurately estimates the white surface beneath the insular cortex, as shown in the Results. The pial surface was estimated using the T2w image to help exclude dura and blood vessels, similar to the HCP pipeline (Glasser et al., 2013). We modified this procedure by applying an optimized value of maximal cortical thickness (=10mm in ‘fake’ space, 5mm in real space like the FreeSurfer default). The surface and volume data in ‘fake’ space was transformed back into the native AC-PC space, and cortical thickness was recalculated in the animals’ real physical space.

The PostFreeSurfer pipeline transformed anatomical volumes and cortical surfaces into the Yerkes19 standard space, performed surface registration using folding information via MSMSulc (Robinson et al., 2014, 2018), generated the mid-thickness surface (by averaging the white and pial surfaces), generated inflated and very inflated surfaces, as well as the myelin map from the T1w/T2w ratio on the mid-thickness surface. The volume to surface mapping of the T1w/T2w ratio was done using a ‘myelin-style’ mapping (Glasser and Van Essen, 2011), in which a cortical ribbon mask and a metric of cortical thickness were used, weighting voxels closer to the midthickness surface. Voxel weighting was done with a gaussian kernel of 2 mm FWHM, corresponding to the mean cortical thickness of macaque (see below). The surface models and data were resampled to a high-resolution 164k mesh (per hemisphere), as well as lower resolution meshes (32k and 10k) for processing diffusion and functional MRI data, respectively.

### Functional Data Processing

Data were motion corrected, corrected for geometric distortions using spin echo field-map correction with TOPUP (Andersson et al., 2003), registered to the structural images using the single-band reference image and BBR (Greve and Fischl, 2009), normalized to grand 4D mean (=10000) and masked (Andersson et al., 2003; Gonzalez-Castillo et al., 2013; Smith et al., 2013). Intensity bias field correction was not done because the functional data were acquired with the pre-scan normalize filter on. The cerebral cortical grey matter voxels were mapped to the surface with the partial-volume weighted ribbon-constrained volume to surface mapping algorithm and voxels having large deviations from the local neighborhood voxels’ coefficient of variation excluded. Data was minimally smoothed at 1.25mm FWHM using geodesic Gaussian surface smoothing algorithm with vertex area correction and resampled according to the folding-based registration (MSMSulc) to a standard mesh in which the vertex numbers correspond to neuroanatomically matched locations across subjects. The subcortical grey matter voxels were processed in the volume using 1.25mm FWHM subcortical parcel-constrained smoothing and resampling. Altogether, these processes transformed the functional data into a standard set of greyordinates (~10,000 [10k] vertices per hemisphere and ~22,000 subcortical voxels) using the Connectivity Informatics Technology Initiative (CIFTI) format (Glasser et al., 2013).

Structured temporal noise arising from imaging artefacts, motion and physiological noise was reduced using a NHP version of multiple-run implementation of FMRIB’s ICA-based X-noisefier (FIX) (“multi-run sICA + FIX”) (Glasser et al., 2018; Griffanti et al., 2017, 2014; Salimi-Khorshidi et al., 2014). Principal component analysis (PCA) was applied to segregate data into structured and unstructured sub-spaces and detect the dimensionality of the structured subspace based on comparison of the data eigenspectrum with a null data eigenspectrum (a Wishart distribution). The structured subspace was decomposed into statistically independent components using spatial ICA and the resulting components were manually classified as “signal” or “noise”, based on their spatial distribution and temporal properties (N=30) (Griffanti et al., 2017, 2014; McKeown et al., 1998). The FIX classifier was then trained on this manual classification and the performance level was characterized in terms of true positive rate (TPR) and true negative rate (TNR). A total of 186 spatio-temporal features were extracted including species-specific vein maps in the standard space and were used for training/classification. The performance of classifier was evaluated by leave-one-out (LOO) accuracy testing for a range of thresholds. The de-noising procedure included linear trend removal, aggressively regressing out 24 movement parameters, which included 6 parameters of rigid transformation, 6 corresponding derivatives and 12 squares of these parameters, and non-aggressively regressing out the noise components (Griffanti et al., 2014). Finally, unstructured noise was attenuated using a Wishart filter (Glasser et al., 2016a) prior to dense connectome analyses.

Information about different categories of fMRI fluctuations were provided by HCP RestingStateStats (Marcus et al., 2013) adapted for monkey. In brief, RestingStateStats quantifies total fMRI variance (prior to any preprocessing) into six categories: high-pass filter, motion, artefacts and nuisance signals (by FIX classification), unstructured noise (by PCA, see above), neural blood oxygenation level dependent (BOLD) fluctuations (by FIX classification), and FIX-cleaned mean global timeseries. The fractional contribution of each category was calculated by dividing by the total fMRI variance.

### Diffusion data processing

Following the HCP pipeline (Sotiropoulos et al., 2013), the diffusion data was normalized for mean intensity of the b=0 volume, corrected for distortion using a spin-echo field-map (i.e. a pair of b=0 volumes acquired in opposite phase), and for eddy-currents and motion using TOPUP and EDDY (Andersson et al., 2003; Andersson and Sotiropoulos, 2016). The images were then registered to the T1w structural image using the undistorted b=0 volume and a 6-DOF boundary-based registration (Greve and Fischl, 2009), transformed into 0.9 mm structural volume AC-PC space (spline interpolation), and masked with a brain mask. The diffusion gradient vectors were rotated according to the rotational information of the rigid transformation matrix from the b=0 to T1w volume. The quality of the diffusion data was assessed using ‘eddyqc’ in FSL (Andersson and Sotiropoulos, 2015; Bastiani et al., 2019). Summary quality metrics consists of SNR calculated for the b=0 images by average intensity divided by standard deviation of b=0 volumes (n=52), and for each b-value with diffusion angular CNR, i.e. the ratio between the standard deviation of the signal predicted by eddy using a Gaussian Process and the standard deviation of the residuals.

Fiber orientation estimation was performed with a model-based parametric deconvolution approach to estimate three crossing fibers per voxel using ‘bedpostx_gpu’ in FSL (Behrens et al., 2007; Hernández et al., 2013; Hernandez-Fernandez et al., 2018) with a burn in period of 3000 and a zeppelin deconvolution kernel (Jbabdi et al., 2012; Sotiropoulos et al., 2016). The uncertainty in the estimated fiber orientations in white matter voxels was compared with the respective uncertainty obtained when using HCP data, for each of three crossing fibers (orientations sorted based on identified volume fraction). Probabilistic tractography was performed on the fiber orientation estimates using FSL’s ‘probtrackx2_gpu’ algorithm (Hernandez-Fernandez et al., 2018) to generate dense diffusion connectomes (Donahue et al., 2016). In brief, we used vertices in the white matter surface and voxels in the subcortical grey matter as a seed of tracking. Streamlines were allowed to propagate within subcortical regions, but they were terminated on exit (Smith et al., 2012). The pial surface and a curvature threshold of 90 degrees were used as stopping criteria. The brain mask calculated in FreeSurfer was used for a waypoint mask through which paths were kept. The step length was set to 0.23 mm, one fourth of voxel size, and the maximum path length to 200 mm. The calculated dense connectomes were created by counting the number of streamlines that terminal on voxels within the seed regions and normalizing by the total number of generated streamlines. These dense connectomes were parcellated using the M132 cortical areas (Markov et al., 2014) to reduce gyral bias and the parcellated connectome matrices were fractionally scaled, symmetrized, and log_10_-transformed (Donahue et al., 2016). The quality of the dMRI data and the success of the tracking algorithm were evaluated with respect to quantitative retrograde tracer data by correlating log_10_-transformed tractography with the corresponding log_10_-scaled fraction of labeled neurons in a source area relative to the total number of label neurons extrinsic to the injected area (Donahue et al., 2016; Markov et al., 2014). Cortico-cortical pathways that did not exhibit connectivity in the retrograde tracer were excluded from the analysis.

Neurite orientation dispersion and density imaging (NODDI) was used to evaluate tissue microstructure associated with neurite composition (a collective term referring to both dendrites and axons) (Zhang et al., 2012). Briefly, NODDI models three compartments (intra-cellular, extra-cellular and CSF) each with different diffusion properties (stick-tensor-ball model), where the diffusion motion in the intra-cellular compartment is assumed to be restricted to within neurites (stick), while that in the extra-cellular compartment is assumed to be a combination of Gaussian anisotropic (tensor) hindered by the presence of neurites, and Gaussian isotropic (ball) in CSF. The model includes two *a priori* assumed parameters of intrinsic axial diffusivity 1.1 μm^2^/ms optimized for grey matter in human (Fukutomi et al., 2018), and isotropic diffusivity 3.0 μm^2^/ms (Zhang et al., 2012), as well as four unknown parameters (intra-cellular volume fraction, concentration parameter of Watson distribution (K), mean orientation of Watson distribution (μ) and isotropic volume fraction (V_iso_). The estimated parameters of orientation dispersion index (ODI) and neurite density index (NDI), as well as diffusion tensor parameters of fractional anisotropy (FA) and mean diffusivity (MD), were mapped onto the cortical surface using an algorithm weighted towards the cortical mid-thickness (Fukutomi et al., 2018).

## Results

### Coil performance

Coil bench tests showed that the unloaded/loaded Q ratio of the individual coil elements were approximately 215/75=2.9. This relatively low Q-ratio results from the small degree of loading and small electromagnetic flux due to the small diameter of the coil elements. Decoupling between adjacent elements was less than −20 dB indicating low mutual inductance between the elements. This produced noise correlation coefficients averaging 0.084 (interquartile range 0.02 and 0.126) with a maximum of 0.395 (see correlation matrix Fig. 2A). High noise correlation was largely constrained to the nearest neighbor elements (see Fig. 1B for element geometry, see also Supplementary Fig. S1 for coil channel-specific noise correlation maps).

**Figure 2.**
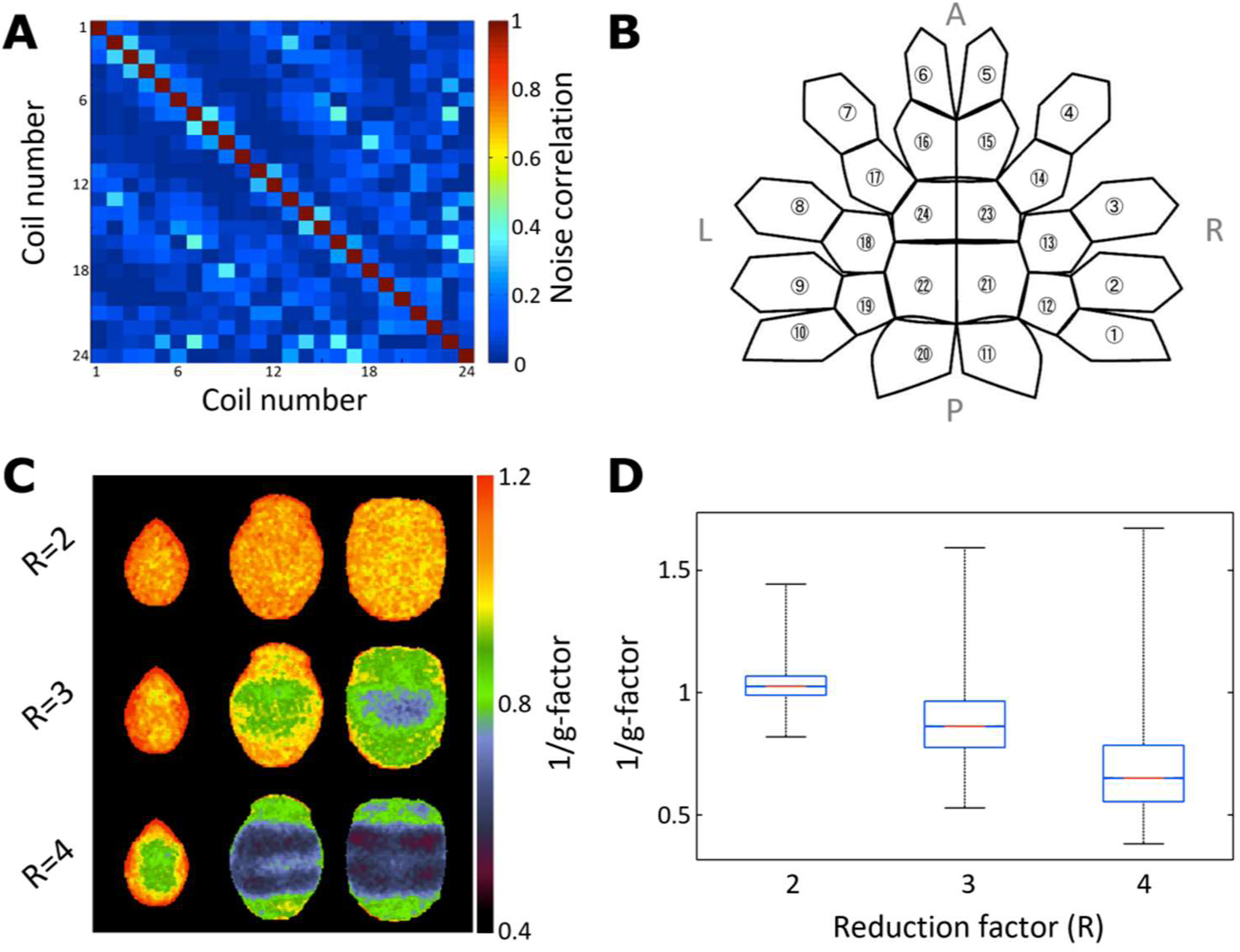
Macaque 24-channel coil performance and geometry. **(A)** Noise correlation matrix. **(B)** Coil element arrangement and labeling flattened into a 2D representation. **(C)** Inverse geometry (1/g)-factor maps using gradient echo imaging with generalized autocalibrating partially parallel acquisitions (GRAPPA) reduction factors (R=2, 3 and 4) in LR-direction used for diffusion MRI (see later). **(D)** The boxplot shows 1/g-factor with respect to reduction factor. While geometric distortions are small with acceleration factor of 2 (1/g=1.03±0.07), further reduction yields large signal degradations. Geometric distortions were evaluated using a phantom whose contour was matched to the average macaque brain.

The inverse g-factor map, a measure of coil element separation, illustrates geometry dependent signal intensity variation due to parallel image reconstruction used for dMRI (Fig. 2C). A reduction factor of two yields an average inverse g-ratio slightly larger than unity (1/g=1.03 ± 0.07; values reported throughout text as mean ± s.d. unless otherwise specified), indicating a small noise cancellation attributable to low element noise correlation and parallel image reconstruction. However, larger reduction factors (R=3 and 4) yield substantial degradation of signal intensity depending on geometry (Fig. 2C, D), suggesting that a maximum GRAPPA of 2 is practical for this coil.

### Macaque Data Quality Evaluation

Structural bias-field corrected T1w and T2w weighted images acquired at 500-μm resolution are shown in Fig. 3A and B for an exemplar single subject. Note the good SNR and contrast of the white matter to grey matter (and to CSF) throughout the brain.

**Figure 3.**
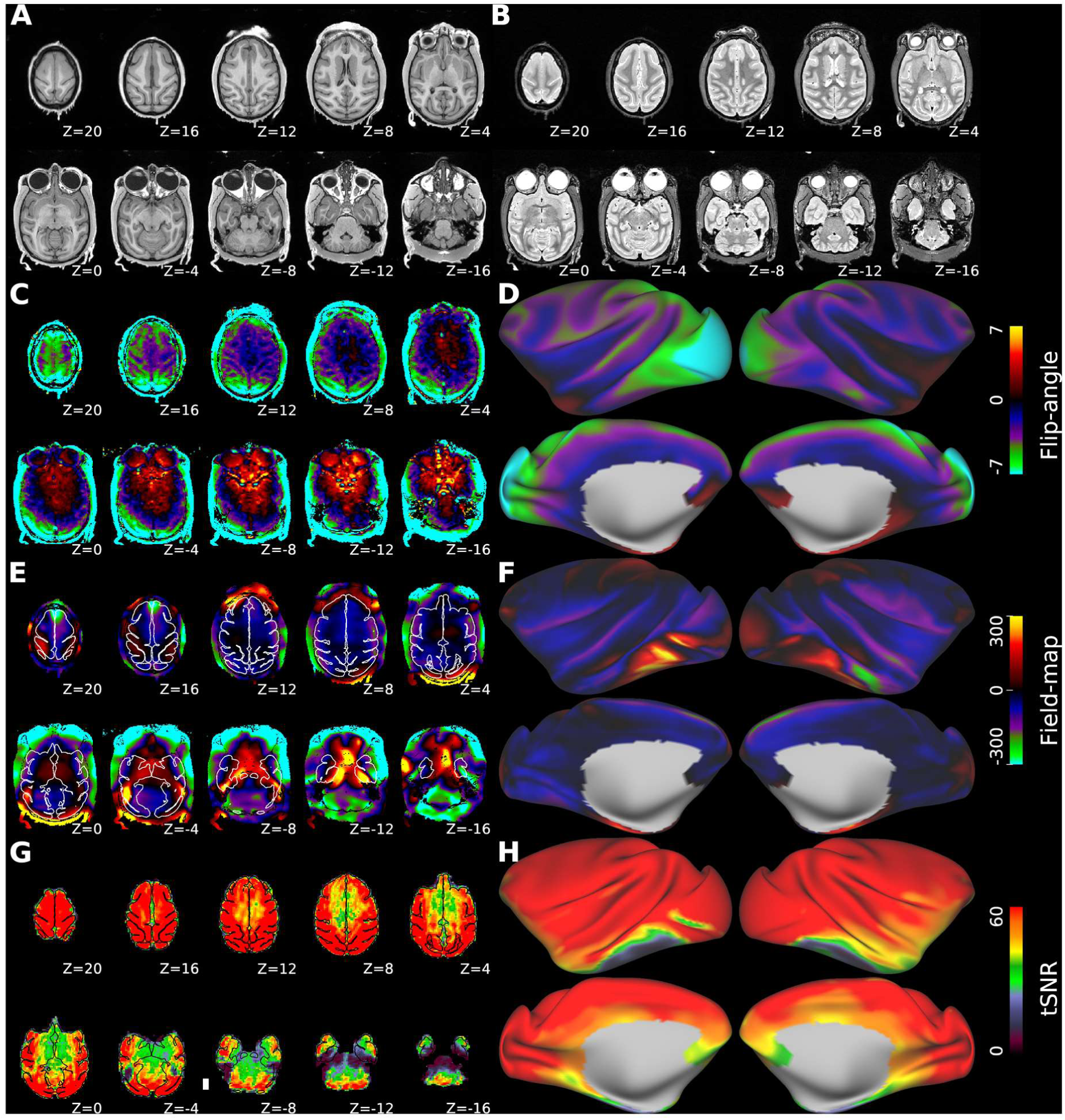
Data quality assessment of structural and functional MRI. Axial slices acquired with 500 μm isotropic resolution T1-weighted MPRAGE and **(B)** T2-weighted SPACE. Flip-angle **(C)** axial and **(D)** surface maps. The values indicate the difference between experimental and nominal flip-angle (90°) in units of degree. B0 **(E)** axial and **(F)** surface field-maps. Unit radian per second. White and black lines (in E and G, respectively) outline the pial surface. Temporal signal-to-noise ratio (tSNR) **(G)** axial and **(H)** surface maps of FIX-cleaned fMRI. The tSNR map was acquired using multiband 2D-EPI sequence (TR=0.755s, TE=30ms, MBF=5, isotropic resolution=1.25mm). Data at https://balsa.wustl.edu/Z44X3

Flip-angle maps indicate that the transmission was slightly higher in subcortical regions compared to cortical structures (Fig. 3C), as expected. However, the surface map (Fig. 3D) indicates that the RF transmission was relatively uniform over the cortical surface (86.6° ± 2.3) (see also Supplementary Fig. S2A for phantom data). Thus, signal intensity and contrast variations at macaque cortical surface attributable to RF-transmission inhomogeneity are modest.

B_0_ volume (Fig. 3E) and surface (Fig. 3F) maps show inhomogeneities, particularly in and near air cavities adjacent to the cerebellum and inferior temporal cortex. These inhomogeneities cause signal intensity loss and spatial distortion in gradient-echo EPI images. Representative tSNR volume (Fig. 3G) and surface (Fig. 3H) maps, acquired with EPI at 1.25 mm isotropic resolution, provide a quantitative estimate for the data quality. The mean FIX-cleaned tSNR in the macaque brain was 51.6 ± 25.6 overall, 67.5 ± 23.7 in the cortical ribbon and 37.3 ± 14.1 in subcortical regions. These macaque tSNR values are higher than the HCP data: the FIX-cleaned tSNR in an exemplar HCP subject was 38.1 ± 15.1 in the whole brain, 43.0 ± 15.2 in the cortical ribbon, and 30.7 ± 10.8 in subcortical regions (Supplementary Fig. S3 and Table S1). However, a relatively low cortical tSNR in lateral occipito-temporal cortex was notable in macaque data (Fig. 3H), which is mainly attributable to a large B_0_ dephasing effect (Fig. 3F).

### Single-Subject Cortical Architecture in Three Macaque Species

FreeSurfer automated segmentation of cortical and subcortical structures using our NHPHCP structural pipeline was reliable across the subjects (Supplementary Fig. S4B), and benefited from additional signal intensity normalization (Supplementary Fig. S4A, see also Supplementary Fig. S2A and S2B for B_1_-transmit and receive fields, respectively). Inspection of pial and white matter surface contours indicates that the automatic segmentation generally followed the contrast boundaries of the T1w image (Supplementary Fig. S4C) and the T2w image (Supplementary Fig. S4D) appropriately, including in challenging thin heavily myelinated regions such as early visual and somatosensory cortex. The subcortical structures including claustrum, pallidum, putamen, were automatically and accurately segmented by the improved subcortical atlas (Supplementary Fig. S4B). The newly added intensity normalization improved the problematic estimation of the white matter surface in the anterior temporal lobe (Supplementary Fig. S5A right), which was not achieved using the default intensity bias field correction (Supplementary Fig. S5A left). The claustrum parcellation strategy also improved the white matter surface just beneath the insular cortex (Supplementary Fig. S5B, right), which often resulted in ‘claustrum invagination’ of the white surface by the default FreeSurfer (Supplementary Fig. S5B left). The claustrum parcellation also improved myelin contrast in the anterior insular area (see next paragraph).

Fig. 4 shows representative cortical surface mapping for three macaque species: Japanese rhesus monkey (*M. fuscata*), rhesus monkey (*M. mulatta*) and crab-eating monkey (*M. fascicularis*), as well as for average of three species (N=12, consisting of N=4 for each species). Although the brain size and surface area were different across species and individuals, we successfully achieved cortical and subcortical parcellation by applying the same Gaussian Classifier Atlas (GCA) and obtained the surface estimation on gyral and sulcal formations (Fig. 4A-H), myelin contrast (Fig.4I-L), which were comparable across species. The total cortical surface area per hemisphere (excluding the non-cortical ‘medial wall’) ranged from 8,093 to 12,897 mm^2^ with an average of 10,052 ± 1,584 mm^2^ (number of hemispheres=24). The average myelin map (Fig. 4L) showed relatively high values in primary visual, sensorimotor and auditory regions and in the “MT+” complex, whereas association areas show relatively low values. These results for myelin maps are in good agreement with each other and with published group average macaque maps (Donahue et al., 2018; Glasser et al., 2014; Glasser and Van Essen, 2011). However, the myelin level in anterior insular cortex tended to be low relative to these earlier maps (Glasser et al., 2014); we consider the present maps likely to be more accurate since this region of agranular insular cortex is very lightly myelinated (Mesulam and Mufson, 1982). Our maps likely benefitted from improved segmentation between claustrum and insular cortex, as described above.

**Figure 4.**
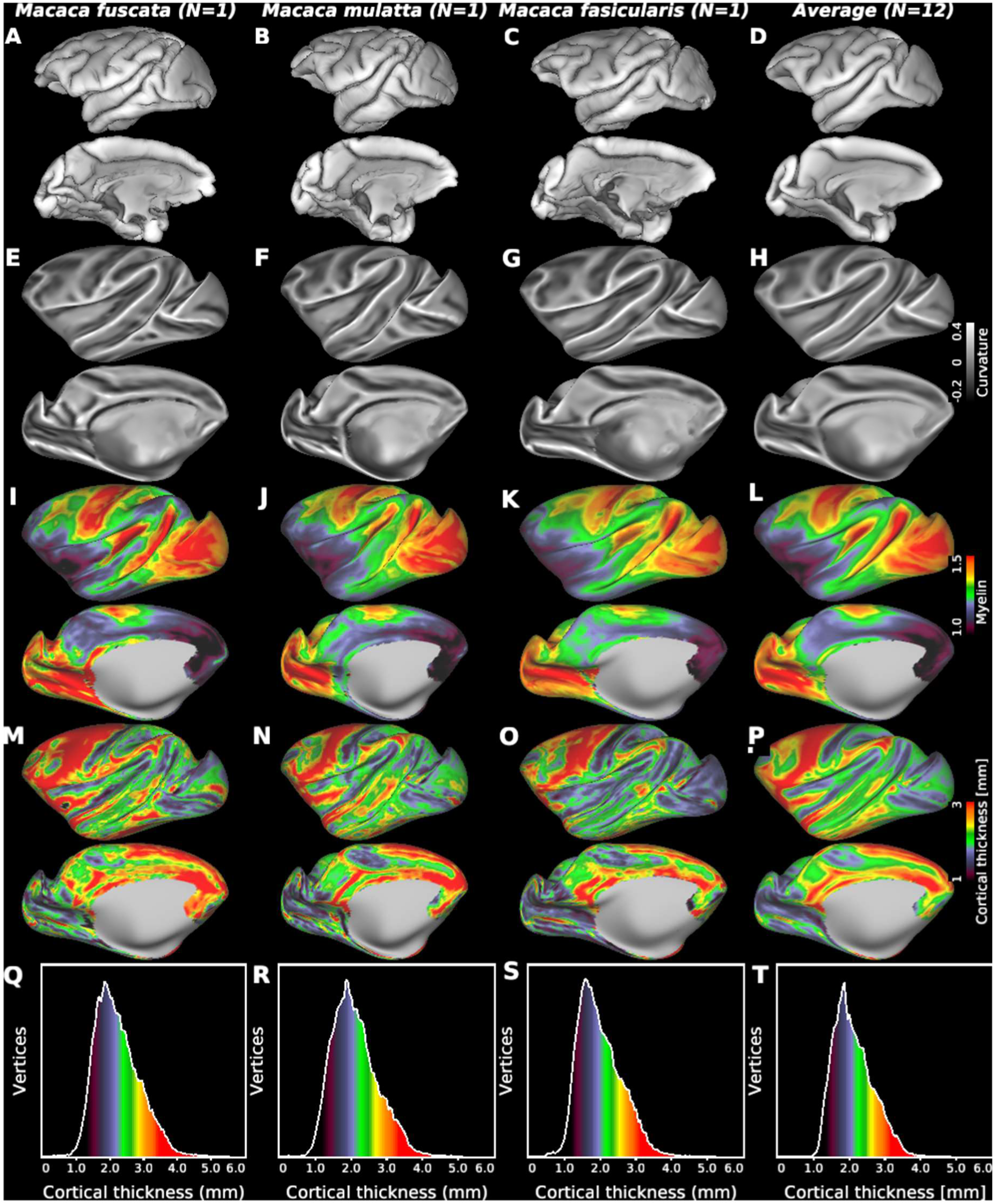
Cortical surface mapping of three widely studied macaque monkeys. Japanese rhesus (*Macaca fuscata*, N=1), rhesus (*Macaca mulatta*, N=1), and crab-eating monkey (*Macaca fascicularis*, N=1) and average maps across the species (N=12; N=4 for each species). **(A, B, C, D)** Pial surface. **(E, F, G, H)** Curvature and **(I, J, K, L)** bias-corrected myelin maps shown on very inflated cortical surface. Cortical thickness **(M, N, O, P)** maps and (**Q, R, S, T**) histograms. Data at https://balsa.wustl.edu/VjjZV

Cortical thickness maps were reasonably consistent across three macaque species (Fig. 4M-P). Most of frontal, anterior insular and temporal cortices are relatively thick, whereas most of visual and parietal cortices are relatively thin. Histograms indicate the distribution of cortical thickness (Fig.4Q-T). Average cortical thickness across species was 2.1 ± 0.54 (median 2.0, N=12). The (lower) fifth percentile of the cortical thickness, evaluated from species average, was 1.38 mm. These estimates indicate that utilizing rfMRI isotropic resolution of 1.25 mm (≈ 2 mm^3^) can capture voxels mainly within the cortical sheet, with modest partial volume effects.

### Data quality of resting-state fMRI

To estimate the optimum multiband factor in fMRI, we determined the relationship between tSNR × sqrt(timepoints) and multiband acceleration factor (Fig. 5A) and found that tSNR x sqrt (timepoints) increases up to a factor of 5, then decreases. This pattern was more evident after the data was processed using ICA-based artefact removal algorithm FIX, which yielded approximately 25% improvement in tSNR. In the cortical ribbon, denoised tSNR x sqrt (timepoints) is clearly highest at MBF=5 (Fig. 5B).

**Figure 5.**
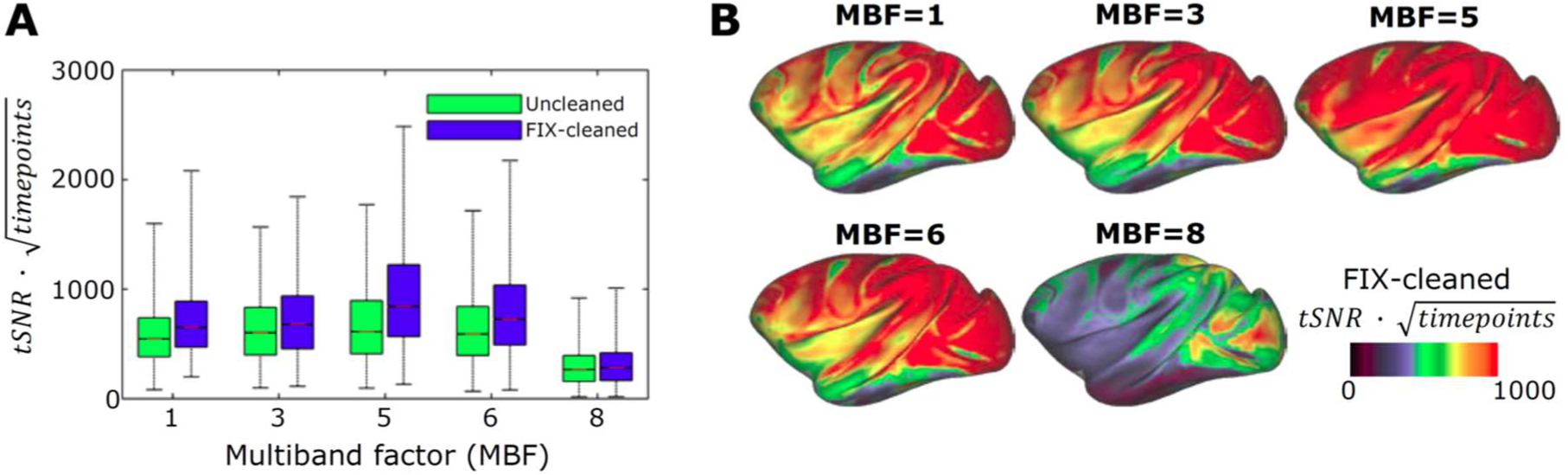
Optimization of fMRI multiband acceleration. **(A)** Relationship between temporal signal-to-noise ratio (tSNR) multiplied by a square-root of acquired time-points and multiband factor (MBF). Acquisition times are matched in the data points (each scan 10 minutes, N=1). The boxplot shows distributions of tSNR in the greyordinates (a total of 26k) for FIX-uncleaned (green) and FIX-cleaned data (blue). **(B)** Cortical surface presentation of FIX cleaned tSNR × sqrt (#timepoints) vs multiband factor. Note that MBF=5 produces the highest tSNR.

The resting-state fMRI runs were analyzed using multi-run sICA + FIX. The resulting sICA components (a total of number of components: 124 ± 29 for each animal, N=30) were manually classified as noise (on average 100 ± 23 components per animal) or signal (24 ± 9 components per animal). The manual classification worked well to train FIX, and the classification accuracy achieved reasonably high performance (Table 1). The LOO accuracy testing showed that mean TPR and TNR ranged between 96.9-99.9% and 95.1-99.6%, respectively, depending on the choice of threshold. A threshold of 20 was used for classification, which resulted in mean TPR and TNR of 99.0% and 98.8%.

**Table 1.** FIX classification accuracy tested by leave-one-out (LOO) in thirty anesthetized macaque data. Abbreviations: TPR=true positive rate of signal components and TNR=true negative rate of true artefact components.

| FIX threshold | 1 | 2 | 5 | 10 | 20 | 30 | 40 | 50 |
| --- | --- | --- | --- | --- | --- | --- | --- | --- |
| TPR (mean) | 99.9 | 99.9 | 99.7 | 99.6 | 99.0 | 98.3 | 97.4 | 96.9 |
| TNR (mean) | 95.1 | 96.1 | 97.4 | 98.2 | 98.8 | 99.0 | 99.3 | 99.6 |
| TPR (median) | 100 | 100 | 100 | 100 | 100 | 100 | 100 | 100 |
| TNR (median) | 96.1 | 96.4 | 97.8 | 98.6 | 98.9 | 99.1 | 99.6 | 100 |

Using RestingStateStats in HCP Pipeline (Glasser et al., 2018; Marcus et al., 2013), the variance in macaque resting-state fMRI runs was divided into six categories. Fig. 6 shows their relative contributions to the total signal variance (38,400 ± 13,000, N=20, see also Table S2). Relative variance estimations in descending order were unstructured noise (70.0 ± 4.8%), high-pass filtered noise (15.3 ± 4.5 %), structured noise (i.e. artefacts and nuisance signals, 6.0 ± 1.5%), (neural) BOLD fluctuations (4.1 ± 2.3%), motion (2.9 ± 1.3%), and FIX-denoised global signal timeseries (1.0 ± 0.7%). In comparison to HCP, unstructured noise accounted for a slightly larger portion in macaque (Fig. 6), which mainly originates from subcortical structures (see Supplementary Fig. S6 for spatial distribution of the variance categories). Furthermore, the relative BOLD contribution was smaller in macaque (4.1%) in comparison to HCP (7.7 ± 2.6%). Taken together, the contrast-to-noise ratio (CNR), defined as ratio between BOLD and unstructured signal, was smaller in macaque (0.21 ± 0.07) than in HCP (0.37 ± 0.08), which may be due to reduced BOLD signals in the anesthetized state (see section Resting-state fMRI in Discussion).

**Figure 6.**
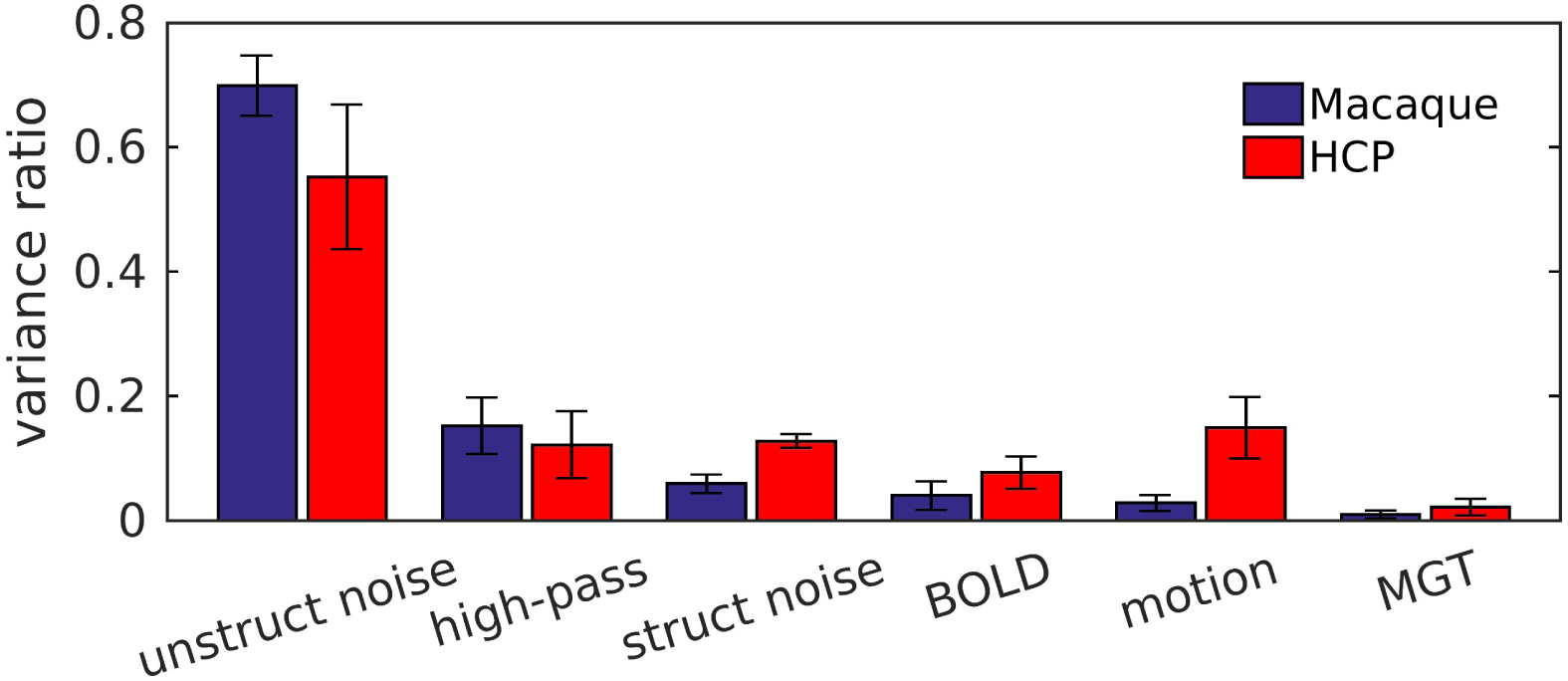
Classification of resting-state fMRI variance and their relative contributions of the total variance in macaque (N=20) and the human connectome project (HCP, N=20). The variances were computed using a development version of the Resting State Stats HCP pipeline. Abbreviations: struct noise=structured noise (scanner artefacts and nuisance signals etc.), BOLD=’neural’ blood oxygen level dependent signal, MGT=FIX-cleaned mean greyordinate timeseries.

One useful way to inspect the data quality is to visualize global (and semiglobal) artefacts in a 2-dimensional heatmap with time on x-axis and parcel (M132) timeseries on y-axis (i.e. greyplot) (Glasser et al., 2018; Power, 2016; Power et al., 2014). Comparison of a representative greyplot prior to any preprocessing (Supplementary Fig. S7A) and after preprocessing (Supplementary Fig. S7B) demonstrated that preprocessing reduced structured artefacts. The mean global timeseries (MGT) also demonstrate that FIX reduced the global signal variance, which in humans is primarily related to respiration after movement artefacts have been removed by sICA+FIX. MGT power spectrum (Supplementary Fig. S7C) revealed distinct peaks within the ventilation frequency range (0.25 to 0.30 Hz). Preprocessing effectively attenuated ventilation artefacts, but only partially attenuated the low frequency, more likely neural, fluctuations (<0.1 Hz). Across subjects, the MGT variance was 2,230 ± 1,530 prior to preprocessing and 170 ± 110 after preprocessing (Supplementary Fig. S7D, N=20). There appears to be relatively less global physiological noise in the macaque data relative to the human data (Glasser et al., 2018; Power, 2016), perhaps because the animals’ respiration was externally controlled by the respirator.

Figure 7 shows a representative resting-state network (RSN) component and seed-based connectivity obtained in a single monkey. Data was from two 51-min fMRI scans, preprocessed for correction of motion, distortion, inhomogeneity, and denoising with multi-run FIX as described earlier. The dense timeseries was further reduced in random noise using Wishart filtering (Glasser et al., 2016a) and was used for seed-based dense connectivity by computing the full correlation. The example RSN component (Fig. 7A) extended positive connectivity over posterior parietal cortex (areas 7A, DP, LIP), precuneus (areas 23, 31), temporo-occipital areas (MST, PGa) and prefrontal cortex (areas 46d, 8b). Temporal properties of this component included low frequency fluctuations, less than 0.2 Hz, which are typical of RSNs. A similar functional connectivity pattern was found using a single greyordinate seed placed over the area 7A (Fig. 7B). Both the RSN signal components (a total of 32 signals) and the dense functional connectome can be interactively viewed in Connectome Workbench after downloading data from the BALSA database (https://balsa.wustl.edu/3ggwG). Overall, these results demonstrate that our experimental setup enables robust functional connectivity detection and analysis.

**Figure 7.**
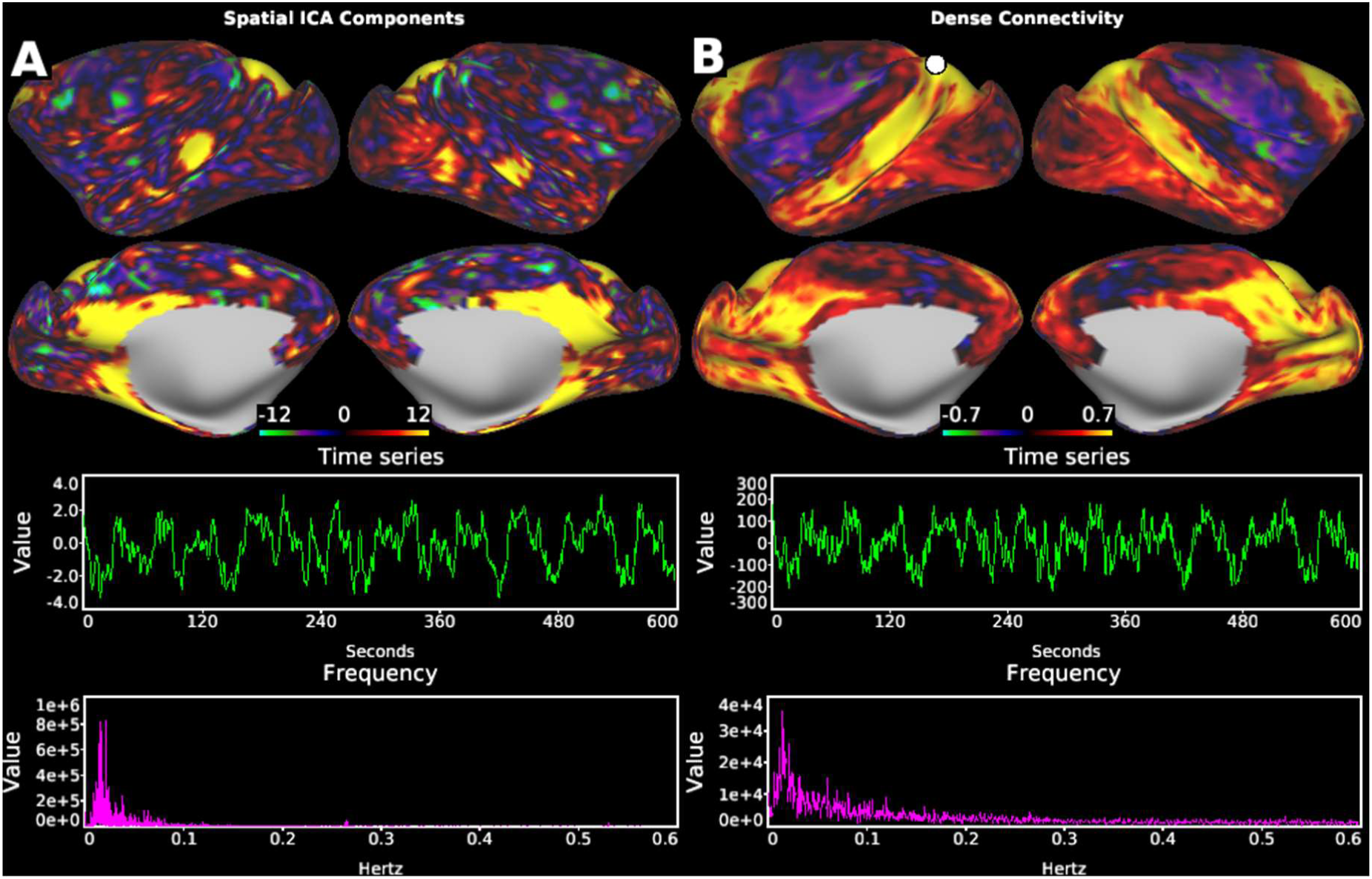
Representative macaque resting-state functional connectivity in a single subject. **(A)** An example resting-state network (RSN) obtained in spatial ICA, which shows positive connectivity over posterior parietal cortex (areas 7A, DP, LIP), precuneus (areas 23, 31), temporo-occipital areas (MST, PGa) and prefrontal cortex (areas 46d, 8b, as defined in M132 atlas). Timeseries and frequency of this component (lower panels) exhibited pronounced low-frequency oscillations. (**B**) Exemplar functional connectivity seeded from a single greyordinate in the area 7A (white circle). Spatial distribution of connectivity resembled to that of the component in (A), as well as timeseries and frequency of the seed signal (lower panels). Data was from two 51-min fMRI scans (subject N=1), preprocessed for correction of motion, distortion, inhomogeneity, and denoising with multi-run FIX. The dense timeseries was further reduced in random noise by Wishart filter and used for seed-based dense connectivity (Pearson’s correlation). Other components classified into signal or noise, and dense connectivity seeded from other vertices can be interactively viewed using Workbench using data at https://balsa.wustl.edu/3ggwG

### Diffusion MRI

Following the HCP paradigm, we used reversed left-right phase-encoding directions in dMRI acquisition to reduce TE, TR and distortion and to increase SNR and angular CNR. An example of image distortion and correction (axial and coronal views) is shown in Supplementary Fig. S8. Image distortions are large near regions with large B_0_ inhomogeneity (i.e. temporal lobe, see Fig. 3E, F). Nonetheless, distortion correction was accurate, albeit with some signal drop-out and degraded SNR in these regions. Mean motion absolute displacement during 30-min acquisition was 0.36 ± 0.07 mm (N=10), ensuring little interaction between head motion, eddy-currents and changes in static magnetic field. In contrast to HCP at 3T (Uğurbil et al., 2013), we used simultaneous MB and GRAPPA acceleration to reduce distortions. Inspection of temporal stability of the dMRI acquisition did not reveal pronounced structural artefacts around the ventricles and basal slices (Supplementary Fig. S9), thus indicating that simultaneous MB and GRAPPA accelerations did not substantially interact with physiological noise (Uğurbil et al., 2013). The dMRI quality assurance measures were similar between this study and the HCP (Fig. 8). Average SNR (whole brain) was 11.6 ± 1.4 in macaque (N=10) and 9.4 ± 0.9 in the HCP (N=10) (Fig. 8A). Exemplar subject data are compared in Supplementary Fig. S10 and Supplementary Table S3. The CNR slightly increased towards higher b-values and was similar across the studies (Fig. 8B). In white matter, three crossing fibers voxels (selected by thresholding at 0.05 of third fiber’s volume fraction) were detected in 59% ± 7% and 57% ± 4% of voxels in macaque and the HCP, respectively (Fig 9D). Finally, the dispersion uncertainties of 1^st^, 2^nd^ and 3^rd^ fiber orientations these voxels exhibited were also similar across the studies (Fig 9E).

**Figure 8.**
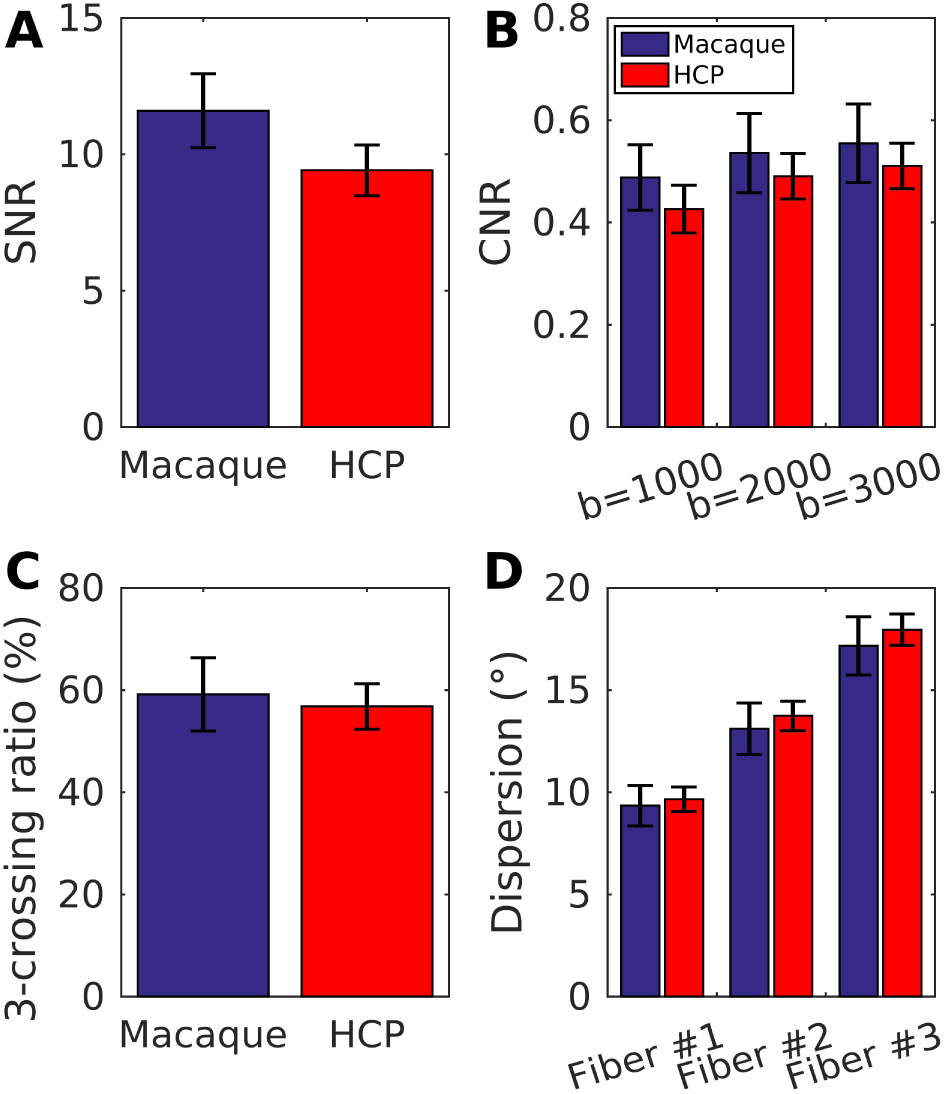
Comparison of dMRI quality measures between macaque and the HCP (blue and red bars, respectively; N=10). Plots show whole brain SNR **(A)** and CNR across b-values 1000, 2000 and 3000 **(B)**, as well as three-crossing fiber ratio **(C)** and dispersion uncertainties (in degree) of 1^st^, 2^nd^ and 3^rd^ fiber orientations in the white matter voxels **(D)**. Overall, the quality measures were comparable across the studies.

**Figure 9.**
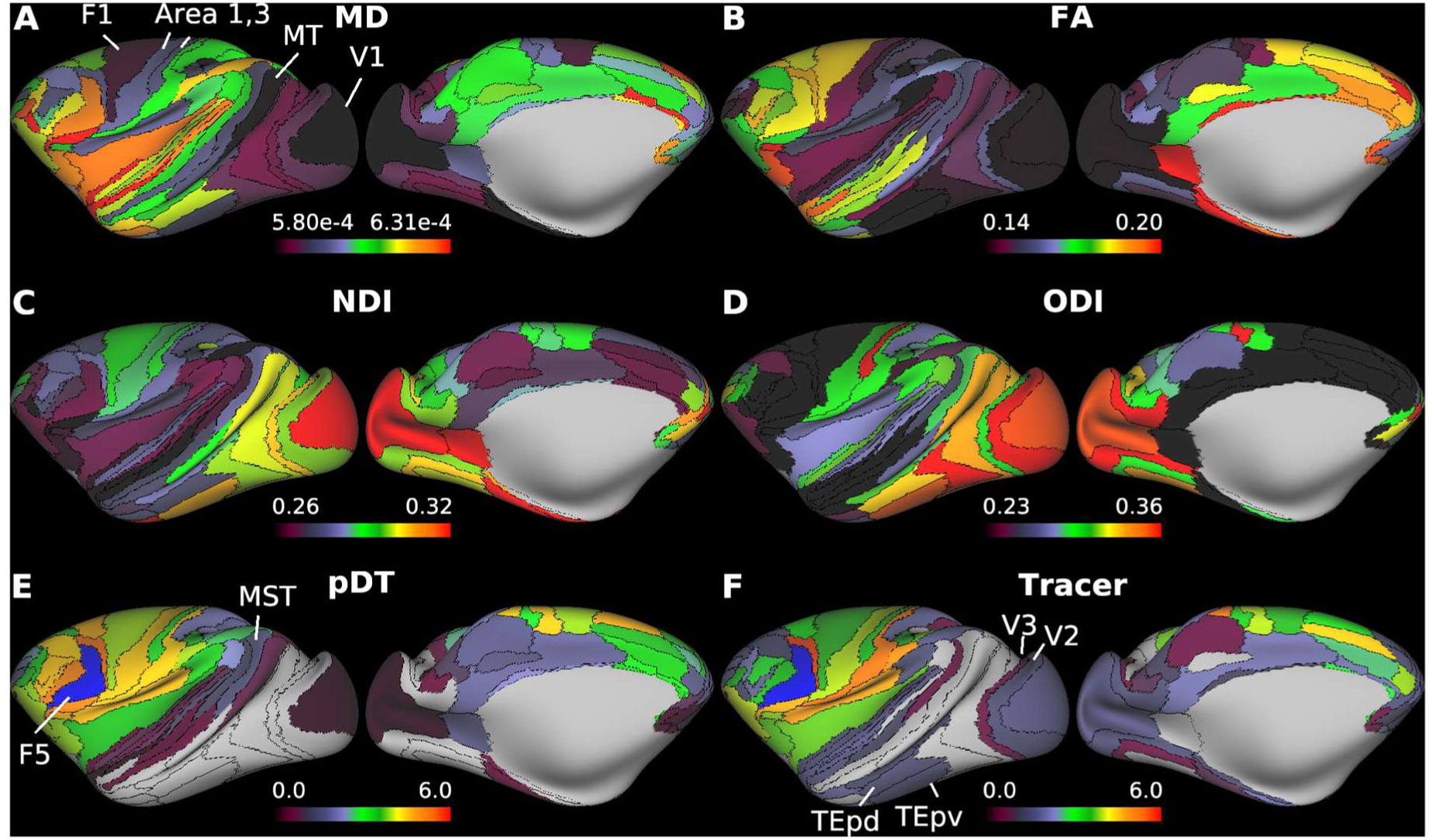
Representative diffusion magnetic resonance imaging (dMRI) applications. Parcellated cortical surface distributions of mean diffusivity (MD) **(A**) and fractional anisotropy (FA) **(B)** calculated in diffusion tensor model, and neurite density index (NDI) and **(C)** orientation dispersion index (ODI) **(D)** calculated in NODDI (see main text; N=6). **(E)** Parcellated diffusion tractography (N=1, ID=A18031601) seed from left premotor area, F5 (blue color) and **(F)** the quantitative ground-truth derived from retrograde tracer injected into F5. Note the correspondence between tractography and tracer connectivities (see main text for details). Data at https://balsa.wustl.edu/zppXg

Figure 9 shows M132 parcellated cortical maps of MD (Fig. 9A), FA (Fig. 9B), NDI (Fig. 9C) and ODI (Fig. 9D) (N=6). The MD is low in the primary motor (F1) and premotor cortices (such as F2, F4, F5), and primary sensory cortices including somatosensory (areas 3, 1, 2), visual (V1) and auditory cortices including core, as well as intraparietal sulcus area (Fig. 9A), whereas the NDI is high in all of these areas. MD and NDI were strongly anti-correlated (R=-0.75, p<0.001). The ODI was high in the periphery of the V1, somatosensory area 1, auditory cortices including core (Fig. 9D) and intermediate in MT and other higher visual areas. The FA was higher in the frontal and anterior temporal cortices and strongly anti-correlated with ODI (R=-0.86, p<0.001). These results are comparable with those observed in the HCP (Fukutomi et al., 2018). The structural connectivity patterns extracted from diffusion tractography (DT) were also parcellated and explored with respect to the published quantitative retrograde tracer data (Fig. 9E, F) (Markov et al., 2014). Comparison between parcellated DT (pDT) seed from area L-F5 and tracer data seeded from area F5 showed a relatively good correlation (R=0.70, p<0.001, for non-zero tracer connections: N=72). However, fidelity of pDT decreased for weak long-distance connections (e.g. false positive connection to MT and MST and false negative connections to V2, V3, TEpd and TEpv), as reported previously (Donahue et al., 2016).

## Discussion

Here, we have presented an adaptation of the HCP’s approach to multimodal MRI acquisition, preprocessing, and analysis to the macaque, using the combination of a custom-made 24-channel receive-coil, high-resolution parallel imaging, and the HCP-NHP preprocessing and analysis pipelines. This approach yields robust estimates of cortical thickness, myelin content, and functional and diffusion measures. Importantly, since the presented protocols used share similar strengths to the HCP image acquisition, and the data is stored in a common geometrical framework system (‘CIFTI greyordinates’), we anticipate that it will facilitate direct multi-modal comparisons with an unprecedented accuracy between macaque and human connectomes. To enable other groups to do HCP-Style analyses in the macaque, this 24-channel macaque coil is available (via Rogue Research; production: Takashima Seisakusho Co. Ltd., Tokyo, Japan) and the data acquisition protocols are freely available from our website (www.nitrc.org/TBA), enabling other investigators to adapt, compare and make the best use of the parallel imaging capabilities of the coil. The HCP-NHP analysis pipelines are also available on github along with the HCP-Style macaque specific FIX training files (https://github.com/Washington-University/NHPPipelines).

### Coil Design

Our multichannel receive coil, fabricated to closely fit a large macaque head (Fig. 1A) will allow routine imaging of macaque monkeys of different species with a range of lateral muscles and head sizes. The close proximity of the coil to the head allows high SNR in the brain with further SNR gains in the cortex produced by the small size of the elements (Fig. 1) (Janssens et al., 2013; Wiggins et al., 2006). This design allowed acquisition of both T1w and T2w structural whole-brain image acquisition with a 0.5mm isotropic resolution in 22 minutes (Fig. 3a, b). In conjunction with homogeneous RF transmission (Fig. 3C, D), these two features enabled automatic and robust subcortical segmentations and reconstructions of pial and white matter surfaces (Supplementary Fig. S4).

Twenty-four receive elements were arranged so as to optimize efficiency of spatial encoding capability in the axial slice direction (Fig. 1B, D). This geometrical arrangement yields a relatively small noise correlation coefficient (0.084), which is smaller than in previous macaque multi-channel coil designs such as 0.12 in a 24-channel (Gilbert et al., 2016) and 0.22 in a 22-channel (Janssens et al., 2013). Our coil design together with slice and in-plane accelerated imaging allowed up to five-fold and two-by-two (MB-by-GRAPPA) accelerations for fMRI and dMRI, respectively. Moreover, this substantially improved the imaging data quality through increased efficiency in accumulation of data volumes (rfMRI: over 8000 volumes; and dMRI: 500 diffusion directions, all acquired in a single session in a period of 140 minutes). Taken together, our 24-channel coil highlights the benefit of accelerated imaging achieved through the geometrical arrangement and low noise correlation of the coil elements.

### Resting-state fMRI

To accurately map BOLD signals onto the cortical sheet, the image resolution (1.25 mm isotropic) was matched with 5^th^ percentile of cortical thickness (Fig. 4N, O, P) to reduce the partial volume effects from white matter and CSF signals (Glasser et al., 2013), following the HCP data acquisition strategy at 3T (resolution 2 mm, the 5^th^ percentile of human cortical thickness (Glasser et al., 2013)). The reduction from an isotropic volume of 2 mm to 1.25 mm, however, incurs a 4-fold SNR penalty. Nonetheless, tSNR of fMRI in macaque (Fig. 3G, H) is superior to that in the HCP acquired with comparable imaging parameters (Supplementary Fig. S3, Table S1). This tSNR gain may be primarily attributed to the close proximity to the animal and small diameter of the receive coil elements, with an additional gain from relatively small bandwidth. This illustrates the power of parallel imaging to overcome a physical size difference of a factor of twelve (macaque and human brain volumes are approximately 100 cm^3^ and 1200 cm^3^, respectively).

While informative, tSNR is not an explicit measure of fMRI sensitivity to blood flow changes induced by neural activity. It is well known that variation of fMRI signal is a mixture of nuisance (e.g. motion and respiration) and neural BOLD components. To obtain insight into the content of our fMRI signals, we categorized different signal sources and found that neural BOLD signal explains approximately 4.1% of the total fMRI variance (in data grand mean scaled to 10,000; corresponding to 773 ± 438 in absolute variance) in anesthetized macaque resting-state (Fig. 6). In HCP fMRI data (awake-state), neural BOLD signal explains approximately 7.7% of total variance (corresponding to 4158 ± 1594 in absolute variance (Glasser et al., 2018; Marcus et al., 2013)). Because the image acquisition protocols and image qualities are similar across the studies (Supplementary Fig. S3), we speculate that the lower BOLD neural signal in our macaque data may be due to, 1) attenuated thalamo-cortical and cortico-cortical synchronization in the anesthetized state, and/or 2) a ceiling effect of signals due to relatively high blood flow, oxygen extraction rate, and saturation in anesthetized macaque brain (Kudomi et al., 2005). This issue may be overcome with widely used contrast agents (i.e. MION) and cerebral blood volume weighted fMRI (Mandeville et al., 1998) to boost CNR. Nonetheless, the relatively small contribution of neural BOLD signal to the total variance highlights the critical importance of post-processing to clean up nuisance signals to obtain functional connectivity estimates that are neurobiologically meaningful. ICA-based FIX denoising has been established to be very successful at removing non-random time-varying spatially specific artefacts (e.g. movement, vascular and cerebrospinal fluid pulsation or scanning artefacts) in the human resting-state fMRI (Griffanti et al., 2017, 2014; Salimi-Khorshidi et al., 2014; Smith et al., 2013). Here, we demonstrated that FIX is also very successful reducing such artefacts (6.0% of total variance, Fig. 6) with over 98% classification accuracy (threshold at 20, Table 1) in the macaque resting-state fMRI. The relative global mean variance and its reduction in macaque (1.5% before cleanup and 1.0% after cleanup) is smaller in comparison to the HCP (3.2% before cleanup and 2.2% after cleanup) (Glasser et al., 2018). This smaller global signal variance in anesthetized macaques can be attributed to more stable global blood flow because respirations and pCO_2_ were regulated by mechanical ventilation (Birn et al., 2006). The majority of the signal variance, however, is unstructured noise (>60%), in particular at subcortical regions that are distant from the coil elements (Supplementary Fig. S6), which can be effectively reduced using parcellation and/or Wishart filtering (Fig. 8B) (Glasser et al., 2016b).

The advantages of our experimental methodology was further demonstrated by the capability to identify an average (across sessions/animals) of 21 ± 9 signal (neural) components at 3T (Fig. 5, for exemplar signal components see Fig. 5 in BALSA). A previous report using group-ICA from six anesthetized macaques at 7T identified 11 RSNs (Hutchison et al., 2011). Our preliminary results replicate several of these RSNs. Taken together, from the data quality perspective, the 24-channel coil yields macaque rfMRI data that can be accurately and sensitively mapped onto cortical sheet and is comparable in quality with the HCP rfMRI data, whereas from the physiology perspective, we must be cautious when making inferences because of the potential effects of anesthesia on both neural activity and neurovascular coupling. We will explore this topic in future work on a specialized coil for awake monkey imaging.

While scaling the fMRI resolution with respect to the cortical thickness is a minimum requirement to accurately localize BOLD signal within the cortical sheet, another important factor is the size of functional imaging voxels relative to the area of the cortical surface for identifying sharp gradient ridges in FC (Glasser et al., 2016a). We found that macaque cortical grey matter surface area is ≈10,100 mm^2^ per hemisphere, which is close to previous estimates of 11,900 mm^2^ (Chaplin et al., 2013) and 9,600 mm^2^ (Donahue et al., 2018). Given that one cortical hemisphere is expected to contain 130-140 cortical areas (Van Essen et al., 2011), an average parcel corresponds to an approximate area of 70 mm^2^ or 70 greyordinates (in our standard 10k greyordinate per hemisphere space for the macaque with 1.25mm average spacing between greyordinates). In comparison, each human cortical hemisphere has an approximate area of 88,200 mm^2^, about is 9-fold larger than in macaque. Since it has 180 cortical areas (Glasser et al., 2016a), an average human cortical parcel corresponds to an area of 490 mm^2^ or ≈160 greyordinates (in the HCP standard 32k greyordinates per hemisphere space for humans). This suggests that identifying clear gradient ridges in FC can be more reliably assessed in the HCP in comparison to our macaque setup, which can be attributed to higher number of greyordinates and 9-fold larger cortical area in humans than in macaques. Viewed from another perspective, since macaque cortex contains 0.85 billion neurons per hemisphere (Herculano-Houzel et al., 2007), a single greyordinate (in 10k space) samples about 85,000 neurons on average. In comparison, since human cortex contain 8.2 billion neurons per hemisphere (Azevedo et al., 2009), so a single HCP greyordinate (in 32k space) samples an average of 270,000 neurons, about three-fold greater than in the macaque. Taken together, while the expected number of greyordinates per cortical area is larger in the human (due to 9-fold larger cortical area of the human brain), our HCP-style approach for the macaque samples fewer neurons per greyordinate (due to the 4-fold smaller voxel volume).

### Diffusion MRI

Spatial resolution is among the most important factors for resolving crossing fiber architecture (Donahue et al., 2016) and microstructural properties such as cortical radial anisotropy (Fan et al., 2017; Sotiropoulos et al., 2016). The ratio between the voxel size and macaque white matter volume for the presented dMRI protocol (0.73 mm^3^ / 23.000 mm^3^ ≈ 3 × 10^-5^) approximately matches 2.5 mm isotropic resolution in the human white matter (16 mm^3^ / 500.000 mm^3^ ≈ 3 × 10^-5^) but is an order of magnitude larger than in the HCP (1.95 mm^3^ / 500.000 mm^3^ ≈ 4 × 10^-6^), although a more precise comparison would require investigations on features such as radii of curvature, tract and blade thickness. Smaller voxel size could aid in distinguishing challenging fiber pathways, however, under our experimental conditions further reduction was impractical due to gradient power and SNR limitations.

To mitigate this limitation, our strategy was to acquire data with exceptionally high angular resolution (500 directions) capitalizing on two-by-two acceleration (out-of-plane MB and in-plane GRAPPA) enabled by the multichannel array coil. The effect of this strategy was shown in the comparable sensitivity to 3^rd^ crossing fibers between species (Fig. 8), despite the resolution limitation in macaque. A recent ex vivo macaque study used high-quality, high-field magnetic field (4.7T), long data acquisition (≈27 h) postmortem and gadolinium enhanced diffusion scans to demonstrate a relatively good correspondence between probabilistic tractography and quantitative retrograde tracer (R=0.55-0.60) (Donahue et al., 2016). Here, we replicated a part of those results (Fig. 8E, F), thus, augmenting the findings of Donahue and colleagues to *in vivo* applications that are within practical time limitations (≈30 min). Taken together, these results suggest that the ‘HCP’-style dMRI data acquisition protocols are well positioned to produce quantitative tractography measures that are neuroanatomically meaningful.

The high spatial resolution with respect to cortical thickness enabled us to carry out cortical surface mapping of neurite properties and to provide preliminary evidence for nonuniformity in the composition and distribution of neurites in macaque cerebral cortex (Fig. 9C, D). Neurite properties are considered important because the density of neurites constitute basic building units (axons and dendrites) of neuronal networks, while ODI provides an indicator of the heterogeneity of neurite fiber orientations, a ratio between tangential and radial fibers (Fukutomi et al., 2018). We found that NDI was highest in V1 and higher than average in other visual representation areas (V2, V3, V4, and MT), somatosensory (1, 2, 3 and A1), motor (M1) and granular prefrontal (Fig. 9C), cortical distributions resembled those of myelin contrast (Fig. 4L). ODI was high in early somatosensory, auditory and visual cortices (Fig. 9D). Together, these results are in good agreement with the HCP data (Fukutomi et al., 2018).

### Towards Improved Macaque Connectomes and Cross-species Connectome Comparisons

The construction of a high-quality connectome requires anatomically accurate definitions of parcels that represent a biologically meaningful partition of brain areas based on their function, architecture, connectivity, and topography (Felleman and Van Essen, 1991; Glasser et al., 2016a; Van Essen and Glasser, 2018). Comparison between transitions in multimodal neuroimaging contrasts, such as MT myelination (Fig. 4C) and functional connectivity (Fig. 8A, E), are particularly suggestive of brain area boundaries (Glasser et al., 2016a). Therefore, the approach to data collection and analysis presented here provides macaque data that may aid in multi-modal parcellation of the macaque and generation of structural and functional connectomes (Glasser et al., 2016b), though a robust delineation of cortical areas into functionally distinct areas will assuredly require analysis of a more extensive dataset.

These HCP-style macaque data also provide an attractive substrate for multi-modal registration across species—in particular, macaques and humans. Just as myelin maps and resting state networks are used to register across human subjects (Robinson et al., 2018, 2014), they could be used to register the cerebral cortex between group averages of humans and macaques. This would allow direct comparisons between human and macaque structural and functional connectivity (Mars et al., 2018b). That said, we expect that the gains for cross-individual registration with areal features in the macaque will be less than those in humans simply because folding patterns and the relationships between folds and areas are less variable in macaques than they are in humans. Additionally, this approach to macaque imaging acquisition and analysis can be used to form structural and functional connectomes in the macaque for comparison with invasively measured tracer datasets (Donahue et al., 2016; Glasser et al., 2016b). Such validation analyses will help to determine the optimal methods for forming structural and functional connectomes in human studies (Jbabdi et al., 2013), where a direct comparison with a gold standard is not available. Future work will also explore cross-species comparisons between macaques and marmosets imaged using specialized hardware and an HCP-style approach. These acquisition and analysis methods can also be applied to study disease models in primate species where controlled and invasive methods can be used to investigate causality and plasticity of structural and functional connectomes and their importance in shaping primate behavior.

### Conclusions

A 24-channel phased-array coil for 3T was constructed and optimized for in vivo parallel imaging of macaque monkey brain. The coil provided high SNR whole-brain coverage and allowed parallel imaging with high speed acquisition by a five-fold and four-fold increase in functional and diffusion MRI, respectively. The data acquisition strategy in combination with the HCP-NHP minimal preprocessing pipelines enabled robust mapping of structural and functional properties onto surface of the cortex. The presented protocols can be acquired within a single imaging session and represent compelling advance in identifying multi-modal cortical topology and structural and functional connectomes in the macaque. Overall, this study demonstrates that MRI studies in animals may benefit from adapting the methodologies introduced by the HCP.

## Notes

Data of figures and supplementary figures are available at https://balsa.wustl.edu/study/show/LPDP Supplementary Information is available in the online version of the paper.

### Acknowledgements

We thank Henry Kennedy for his helpful comments, and Stephen Frey, Nobuyoshi Tanki, Chiho Takeda, Akihiro Kawasaki, Reiko Kobayashi, Kenji Mitsui and Hanako Hirose for their technical help. This research is partially supported by the program for Brain/MINDS and Brain/MINDS-beyond from Japan Agency for Medical Research and development, AMED (JP18dm0207001, JP18dm0307006), by RIKEN Compass to Healthy Life Research Complex Program from Japan Science and Technology Agency, JST, by MEXT KAKENHI Grant (16H03300, 16H03306, 16H01626, 15K12779 to T.H. and JP26640065, JP16H06531 to Y.G.), NIH F30 MH097312 (M.F.G.) and RO1 MH-60974 (D.C.V.E.) and by the National Institutes of Natural Sciences (NINS) program for cross-disciplinary study (2013-2015) (Y.G.) and by Wellcome Trust (SMS, JS). The author Yoshihiko Kawabata holds financial conflict of interest because the coil is produced by his company (Takashima Seisakusho Co. Ltd., Tokyo, Japan) and distributed through his collaboration company (Rogue Research, Montreal, Canada). The other authors have no conflicts of interest to declare.

## Author Contributions

T.H., and M.F.G. designed the study.

J.A.A, Y.H., M.F.G. and T.H. analyzed data.

M.F.G., C.J.D., and T.H. developed new software tools.

J.A.A, T. O., M. O. Y. K, Y.U, K.M, K. N., M.Y, A.Y., Y.G. and T.H. performed experiments.

J.A.A, M.B., M.F.G, T.S.C, S.J., S.N.S, S.S, D.C.V.E and T.H. wrote the paper.

