## SupplementalMaterial for "Towards HCP-Style Macaque Connectomes: 24-Channel 3T Multi-Array Coil, MRI Sequences and Preprocessing"

**Supplementary Table S1.** Comparison between macaque and the human connectome project (HCP) resting-state fMRI temporal signal-to-noise ratio (tSNR). Mean (s.d.).

| | Data type | Whole brain | Cortical ribbon | Subcortical areas |
|---|---|---|---|---|
| **Macaque (ID Soy)** | Uncleaned | 45.9 (21.5) | 59.9 (19.5) | 34.2 (12.1) |
| | FIX cleaned | 51.6 (25.6) | 67.5(23.7) | 37.3 (14.1) |
| **HCP (ID 100307)** | Uncleaned | 30.8 (11.8) | 33.6 (11.8) | 23.9 (8.3) |
| | FIX cleaned | 38.1 (15.1) | 43.0(15.2) | 30.7(10.8) |

**Supplementary Table S2.** Comparison between macaque and the human connectome project (HCP) resting-state fMRI categorized absolute variance. The mean variances (s.d.) were computed using a development version of the Resting State Stats HCP pipeline (N=20). Shared variances are not shown. Note that the total absolute variance is over two-fold larger in HCP.

| | Total | Unstructured noise | High-pass | Structured noise | BOLD | Motion | Cleaned MGT |
|---|---|---|---|---|---|---|---|
| **Macaque** | 38,356 (13,028) | 28,316 (9,589) | 5,393 (2,976) | 2,575 (819) | 773 (438) | 794 (351) | 171 (114) |
| **HCP** | 82,599 (24,630) | 41,037 (10,833) | 14,115 (11,247) | 10,920 (4,936) | 4,158 (1,594) | 13,179 (6,880) | 1,182 (831) |

**Supplementary Table S3.** Comparison between macaque and the human connectome project (HCP) diffusion MRI (b=0 s/mm$^2$) temporal signal-to-noise ratio (tSNR). Mean (s.d.). Note that 'eddy_combine' was not applied to the HCP data for averaging volumes across different phases.

| | Whole brain | Cortical ribbon |
|---|---|---|
| **Macaque (ID Cranky)** | 12.7 (6.2) | 17.0 (3.7) |
| **HCP (ID 100307)** | 11.3 (4.2) | 13.2 (2.7) |

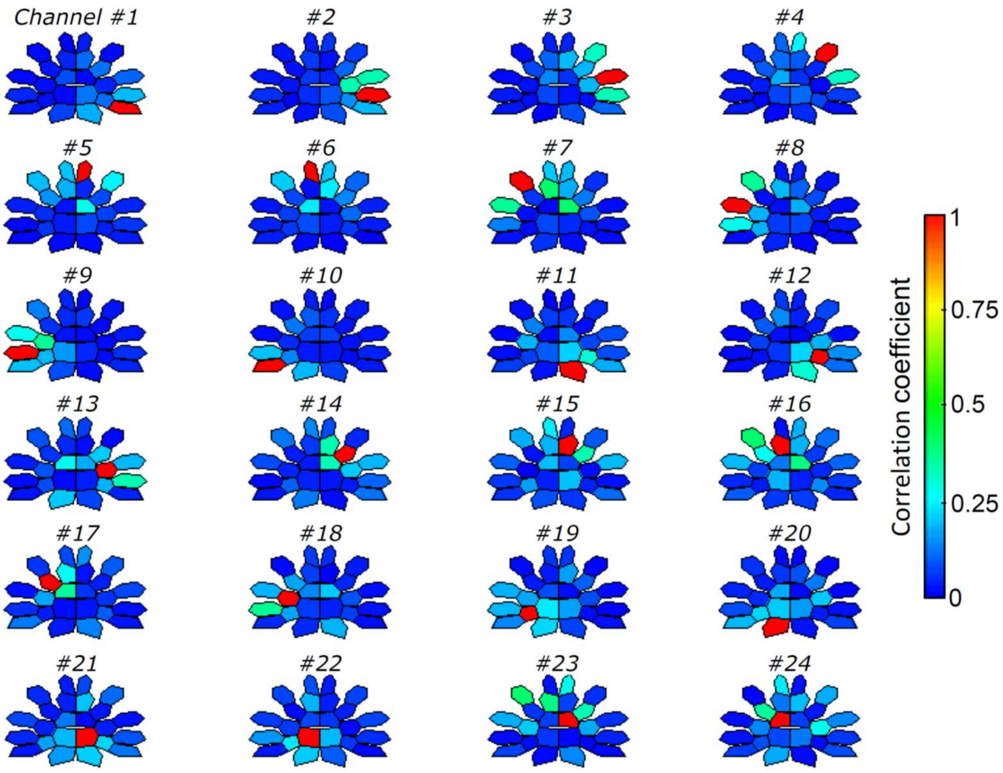


**Supplementary Figure S1**. Coil channel specific noise correlation maps.
The colorbar indicates correlation coefficient. Each seeded channel is indicated by the red color. Noise coupling is mostly limited to next-neighbor interaction and is symmetrical across anterior-posterior (top-down) direction.

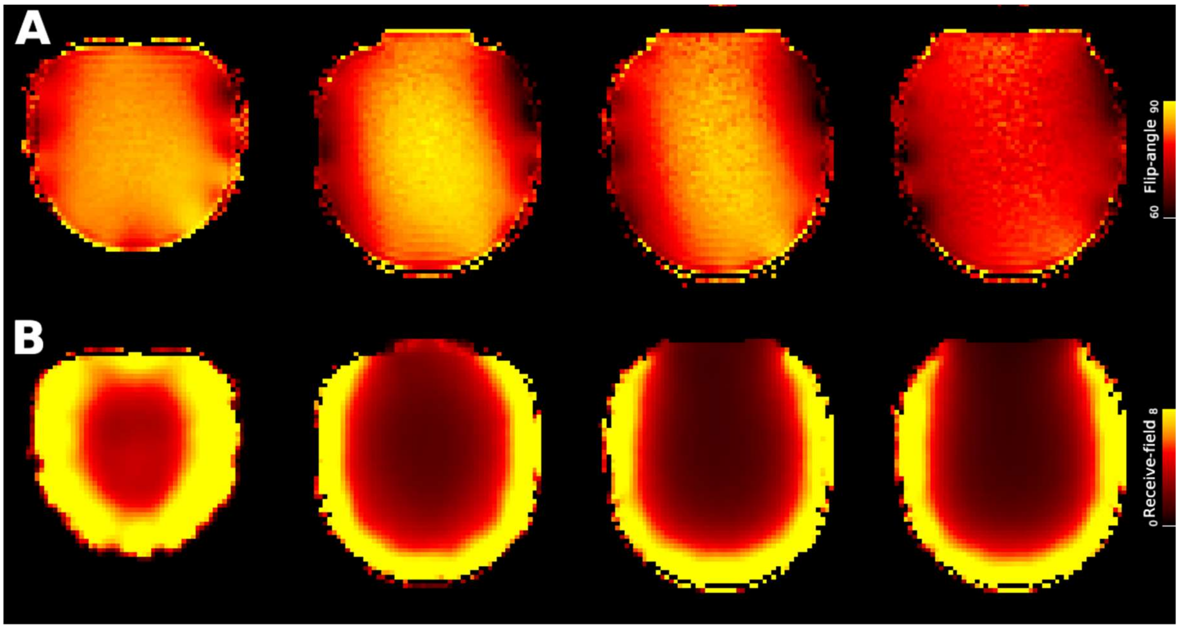


**Supplementary Figure S2.** Biasfield over a phantom, which matched the inner contour of the coil. $B_1$ **(A)** transmission (colorbar indicates flip-angle) and **(B)** receive fields. Data at https://balsa.wustl.edu/Mx93w

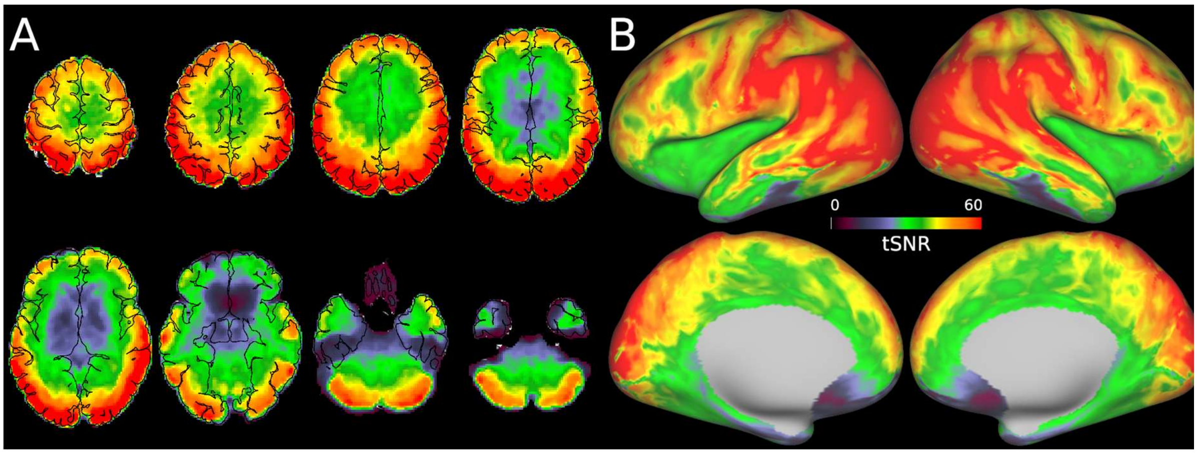


**Supplementary Figure S3**. The human connectome project (HCP) FIX-denoised resting-state fMRI data temporal signal-to-noise ratio (tSNR). Representative **(A)** axial and **(B)** surface maps for a single human subject (ID: 100307 in the HCP). The tSNR map was acquired using 32-channel coil and 2D gradient-echo EPI (TR=720ms, TE=32ms, Multi-band factor=8 and isotropic resolution=2mm (Smith et al., 2013)). Data at https://balsa.wustl.edu/B4gPl

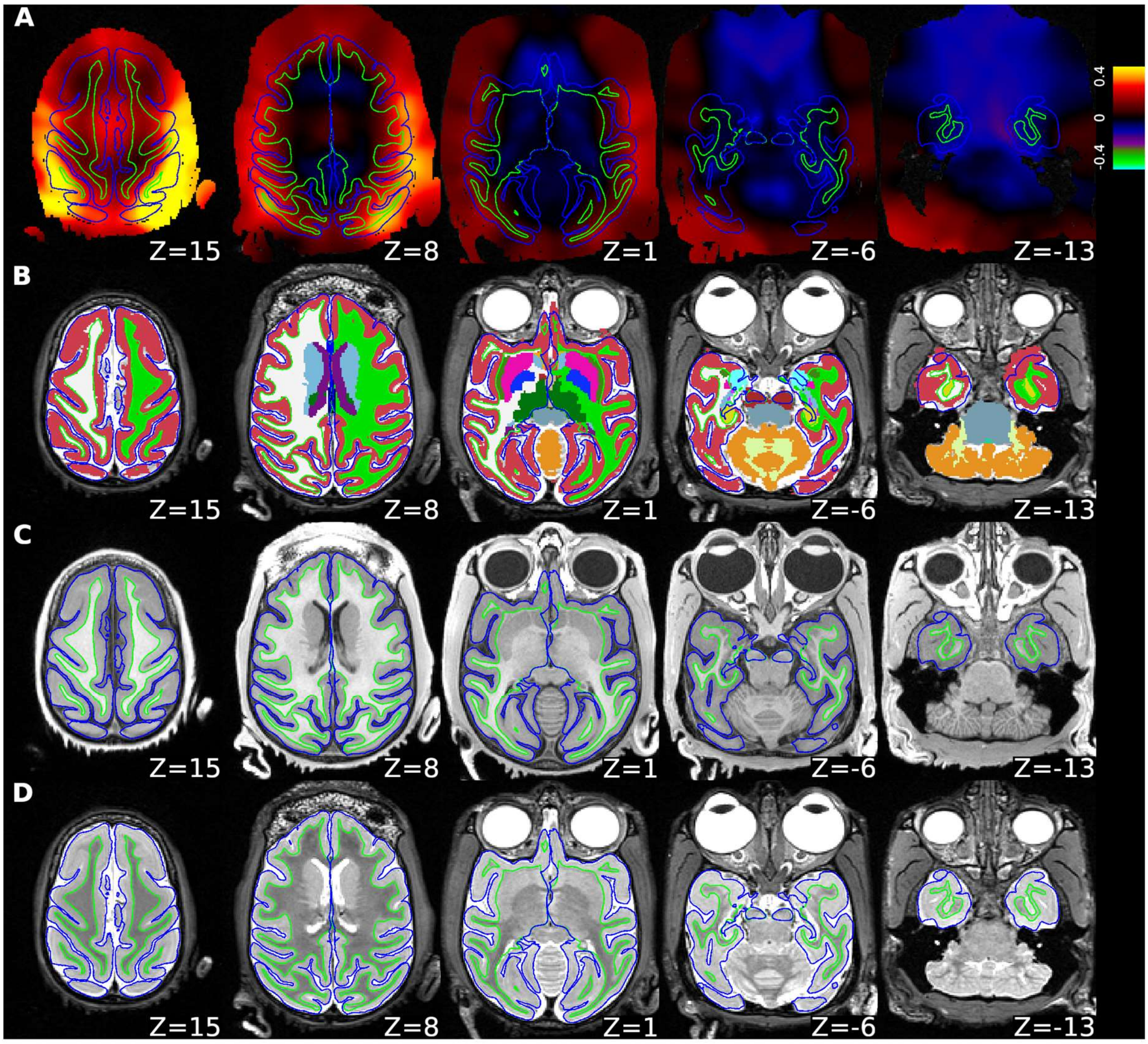


**Supplementary Figure S4**. Biasfield, segmentation and structural images of an exemplar macaque. **(A)** $B_1$ bias-field estimated using the signal intensity ratio between T1w and T2w images. The colorbar indicate the bias-field in log units. **(B)** Brain segmentation. Note that segmentation of claustrum was achieved. **(C)** T1w and **(D)** T2w images. Reconstructed cortical surfaces for white matter (green line) and pial (blue line) are overlaid on all of images. Data at https://balsa.wustl.edu/lL7DL

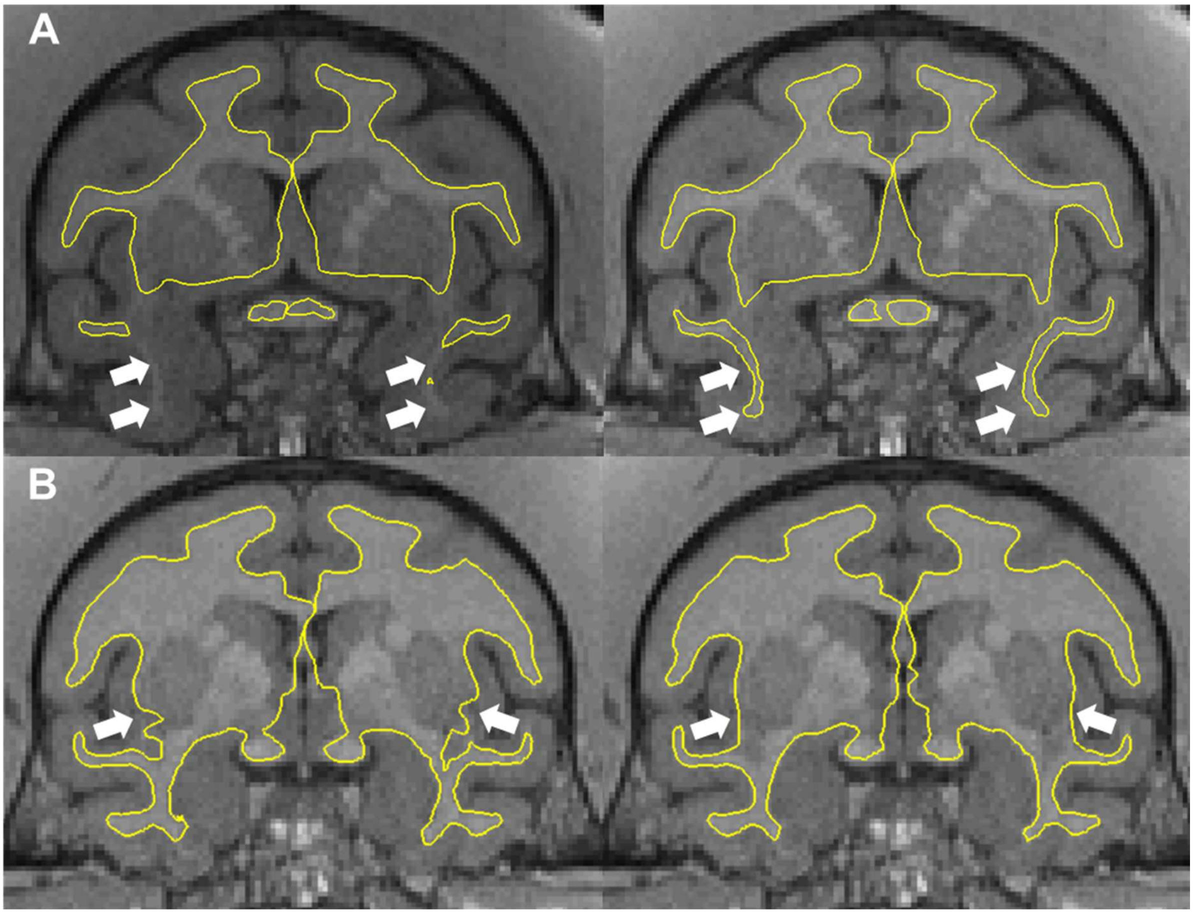


**Supplementary Figure S5**. Improved estimation of white surface.
**(A)** 'Anterior temporal lobe problem', in which white matter blade in the anterior temporal lobe (white arrows) was missing in the default FreeSurfer pipeline (left), was improved in the newer version of the HCP-NHP pipeline. This was achieved by including an additional process for normalizing the intensity for T1w images. Note that the contrast of T1w images (grey color) was slightly increased after the normalization (right) as compared to that before (left). **(B)** 'Claustrum invagination problem', in which white matter surface was invaginated into the claustrum in the default FreeSurfer pipeline (white arrows), was improved by enabling an automated claustrum segmentation followed by white matter surface estimation. Yellow lines indicate white matter surfaces.

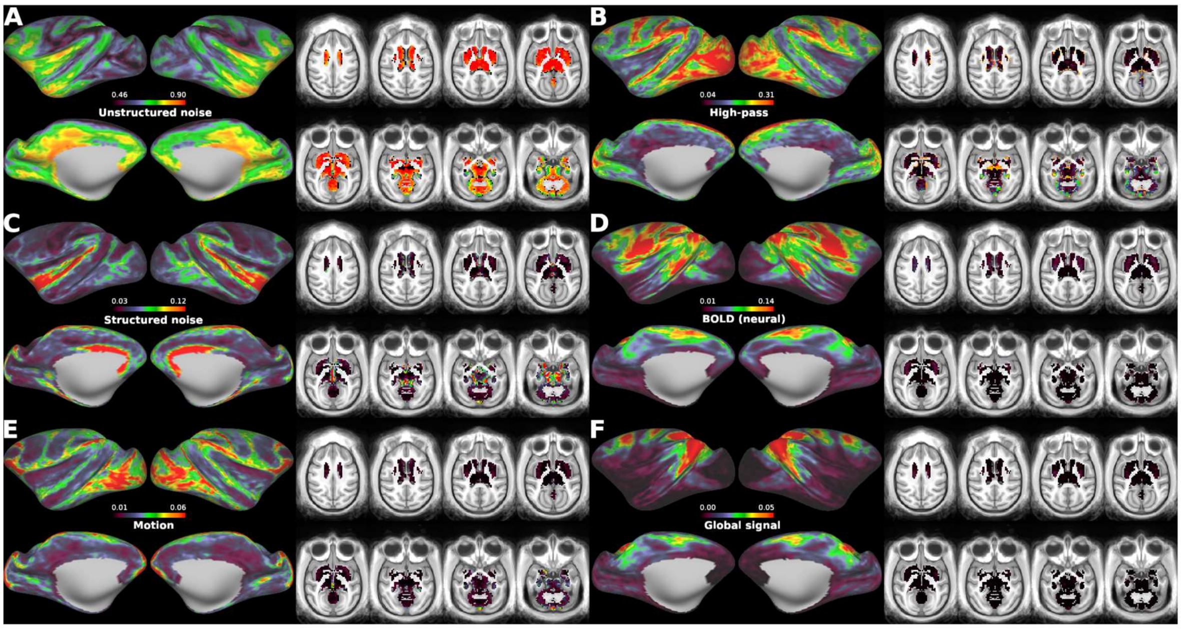


**Supplementary Figure S6.** Distribution of relative signal variance in macaque resting-state fMRI. **(A)** Unstructured noise, **(B)** high-pass filter, **(C)** artefacts and nuisance signals (i.e. structured noise), **(D)** blood oxygen level dependent (BOLD), **(E)** motion and **(F)** FIX-cleaned mean global grey matter signal. The relative variances were computed using a RestingStateStats algorithm in the human connectome project (HCP) pipeline (Glasser et al., 2018) and averaged across subjects (N=20). Data at https://balsa.wustl.edu/qNxgx

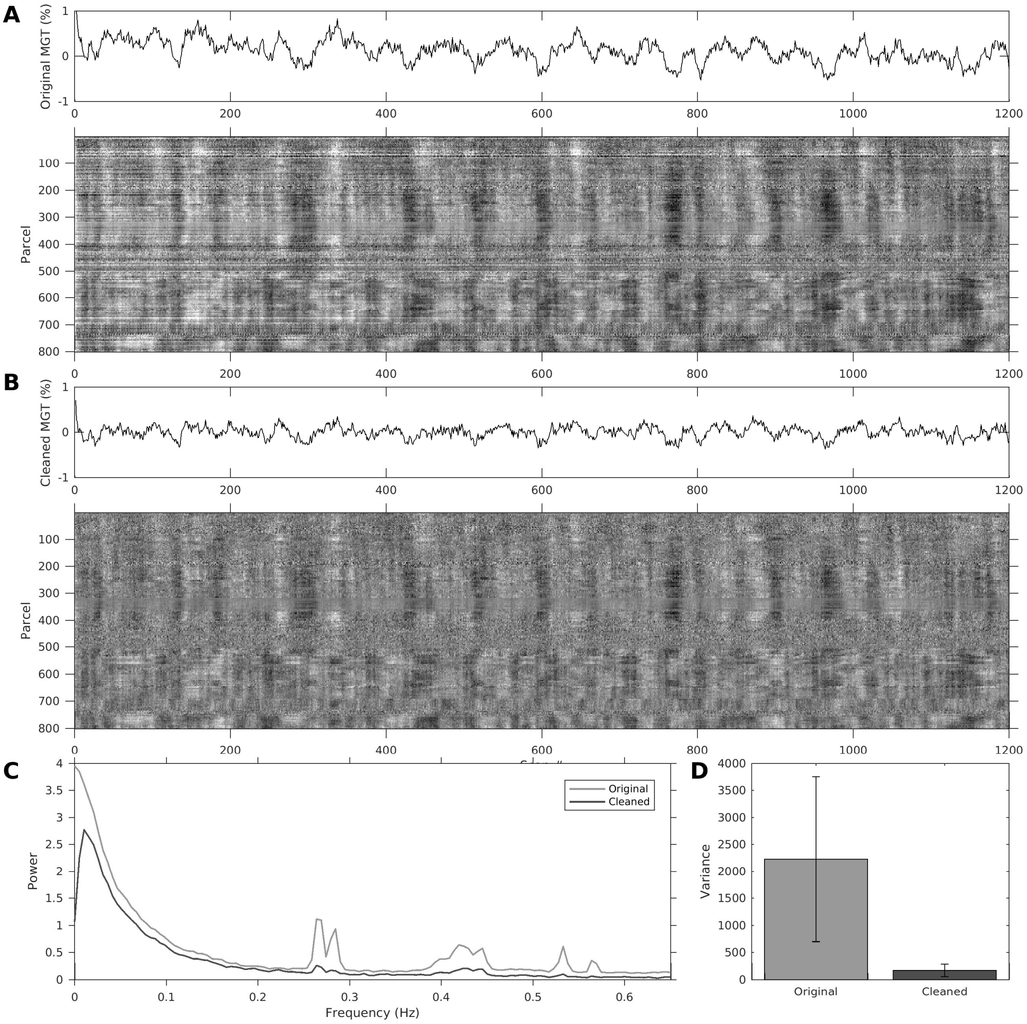


**Supplementary Figure S7**. Mean global timeseries (MGT) and greyplot of **(A)** uncleaned (without any preprocessing) and **(B)** cleaned (motion correction, band-pass, FIX-cleaning) data in a single representative subject. The greyplots are scaled according to % parcel mean signal (±2%) balanced according to parcel size and ordered by hierarchical clustering (Ward's method) for visualization. **(C)** Power spectrum of MGT. Note that FIX attenuated ventilation artefacts at 0.25-0.30 Hz. **(D)** MGT variance was substantially reduced by preprocessing (including band-pass filter, motion correction and FIX-cleanup, N=20).

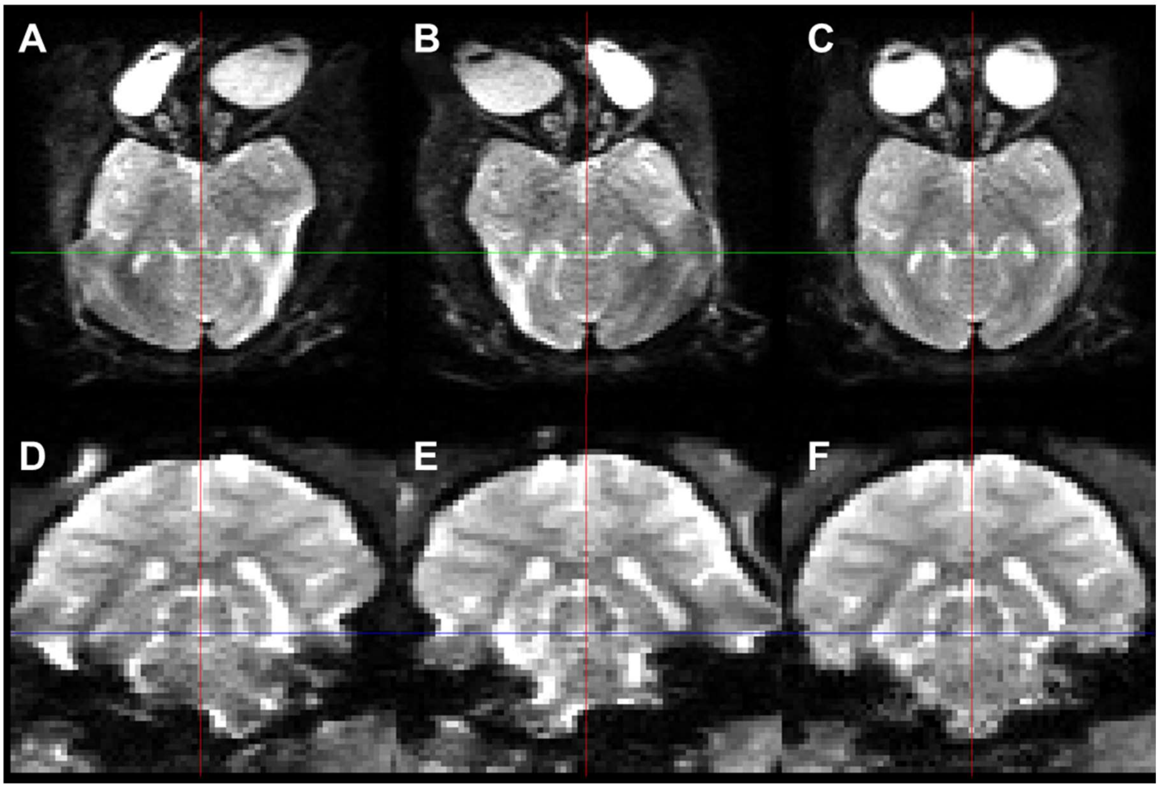


**Supplementary Figure S8.** Correction of susceptibility distortions using reversed phase encoding (PE) b=0 s/mm$^2$ images.
**(A, D)** Left-right and **(B, E)** right-left PE directions. **(C, F**) Distortion corrected image. Axial (top) and coronal (bottom) views.

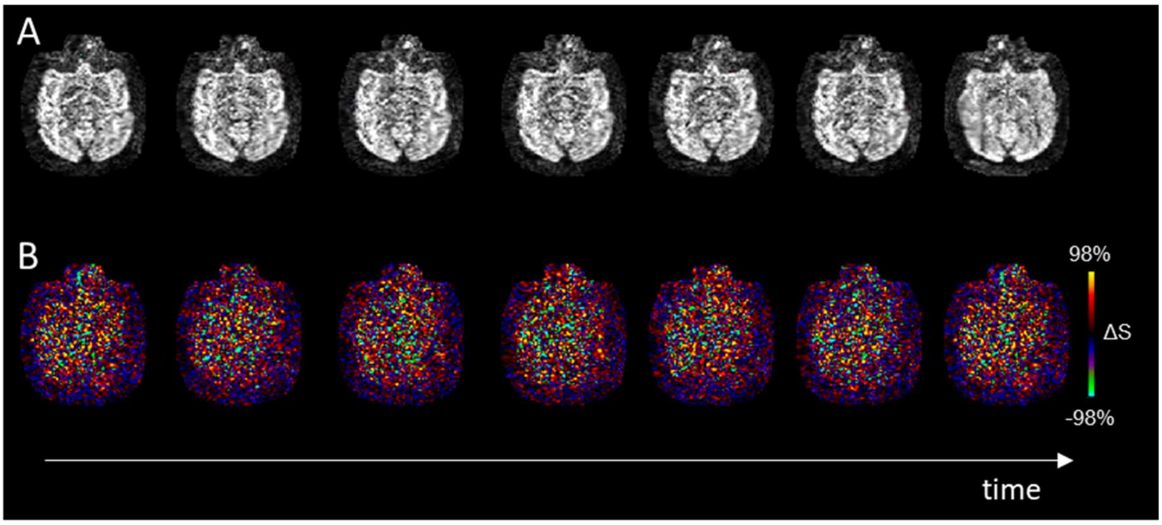


**Supplementary Figure S9.** Temporal stability of diffusion MRI with slice (MBF=2) and in-plane acceleration (GRAPPA=2).
**(A)** Consecutive b=1000 s/mm$^2$ images with same diffusion-weighting direction. **(B)** Corresponding images with mean signal intensity across frames subtracted (i.e. $\Delta S=S-\text{mean}(S)$). We found no strong interaction between diffusion-weighting, accelerated imaging and physiological pulsations.

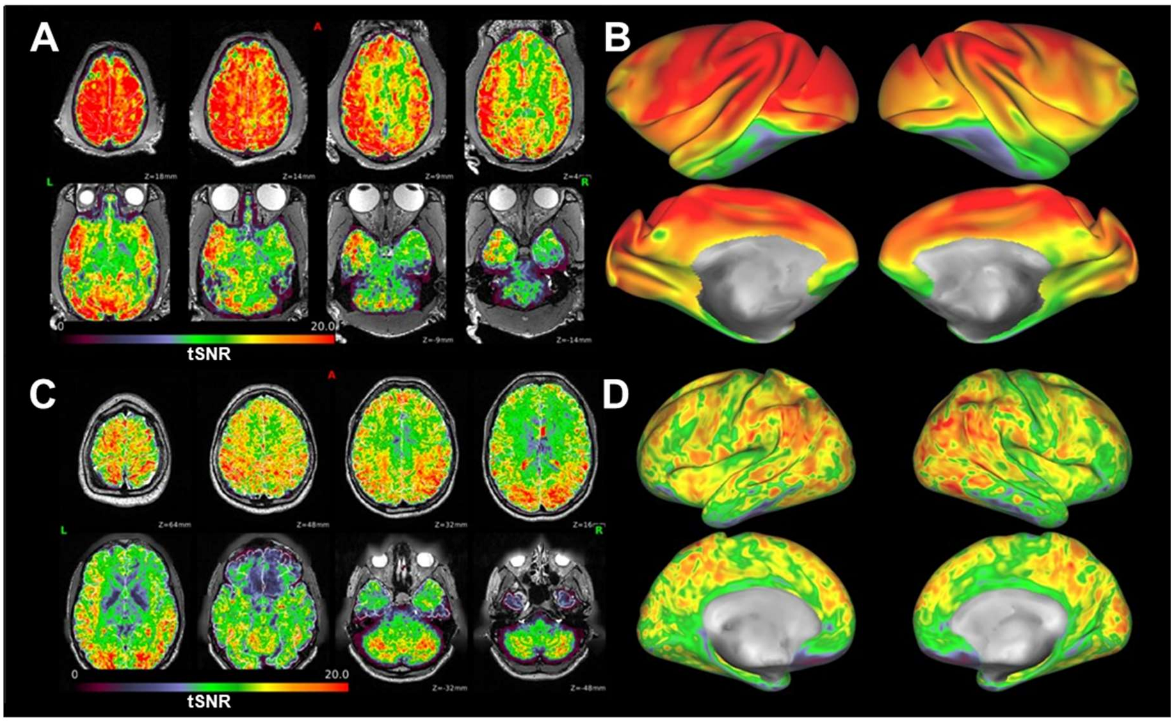


**Supplementary Figure S10**. Comparison between macaque and HCP diffusion (b=0 s/mm$^2$) temporal signal-to-noise ratio (tSNR).

Representative **(A)** axial and **(B)** surface macaque tSNR (species: Macaca Mulatta, body weight=4 kg) acquired with a 3T scanner. Representative human **(C)** axial **(D)** surface tSNR from the Human Connectome project (HCP) (ID 100307) acquired with a 3T connectome scanner. The macaque data was obtained with 24-channel coil and 2D spin-echo EPI sequence with isotropic resolution=0.9 mm, TR=3400 ms, TE=73 ms, readout bandwidth=1086 Hz/pixel, Multiband factor (MBF)=2 and GRAPPA=2, while human HCP data was obtained with 32-channel coil and 2D spin-echo EPI with isotropic resolution=1.25 mm, TR=5520 ms, TE=90 ms, readout bandwidth=1488 Hz/pixel and MBF=3 (Sotiropoulos et al., 2013). Note that macaque acceleration was two-by-two (MBF=2 and GRAPPA=2) whereas in HCP acceleration was three-by-one (MBF=3 and no GRAPPA).
